# RNAseq analysis of rodent spaceflight experiments is confounded by sample collection techniques

**DOI:** 10.1101/2020.07.18.209775

**Authors:** San-Huei Lai Polo, Amanda M. Saravia-Butler, Valery Boyko, Marie T. Dinh, Yi-Chun Chen, Homer Fogle, Sigrid S. Reinsch, Shayoni Ray, Kaushik Chakravarty, Oana Marcu, Rick B. Chen, Sylvain V. Costes, Jonathan M. Galazka

## Abstract

To understand the physiological changes that occur in response to spaceflight, mice are transported to the International Space Station (ISS) and housed for variable periods of time before euthanasia on-orbit or return to Earth. Sample collection under such difficult conditions introduces confounding factors that need to be identified and addressed. We found large changes in the transcriptome of mouse tissues dissected and preserved on-orbit compared to tissues from mice euthanized on-orbit, preserved, and dissected after return to Earth. Changes due to preservation method eclipsed those between flight and ground samples making it difficult to identify spaceflight-specific changes. Follow-on experiments to interrogate the roles of euthanasia methods, tissue and carcass preservation protocols, and library preparation methods suggested that differences due to preservation protocols are exacerbated when coupled with polyA selection. This has important implications for the interpretation of existing datasets and the design of future experiments.

## Introduction

Spaceflight places multiple stresses upon the human body including altered gravity fields and exposure to cosmic radiation, which lead to health risks for spacefaring humans (Institute of Medicine, 2008). Decades of research on astronauts has begun to reveal how humans respond to the spaceflight environment (Garrett-Bakelman et al., 2019) but physiological monitoring of astronauts is still limited. Thus rodent models have been essential for advancing our understanding of how mammals — including humans — respond to spaceflight. This includes the impact of spaceflight on muscle structure (Shen et al., 2017; Spatz et al., 2017; Tascher et al., 2017), liver (Beheshti et al., 2019; Jonscher et al., 2016) and immune functions (Pecaut et al., 2017; Rettig et al., 2017; Ward et al., 2018).

Despite success of the rodent model, sample collection under such difficult conditions introduces confounding factors that need to be identified and addressed. These are related to hardware limitations, small sample size, and severe restraints on astronaut crew availability. Successful experiments must work within these constraints to produce meaningful insights. In response, the first Rodent Research (RR) mission established new capabilities for conducting reliable long-duration experiments using rodents with on-orbit sample collection. Animals can either be euthanized onboard the ISS by or returned to Earth alive. Both approaches introduce confounding factors. The former is experimentally challenging but preserves the sample during exposure to microgravity, while the latter exposes the animal to re-entry stresses and sampling occurs only after a variable lag between landing and euthanasia — essentially sampling re-adaptation to Earth conditions in addition to the response to spaceflight. Inconsistent handling of samples necessitates a clear understanding of how dissection and preservation protocols affect downstream data generation.

We previously showed, using transcriptomic, proteomic and immunohistochemical data from the Rodent Research-1 (RR-1), Rodent Research-3 (RR-3) and Space Transportation System (STS)-135 missions, that lipotoxic pathways are activated in rodent liver in two different strains of mice that were flown for as long as 42 days in space (Beheshti et al., 2019). Because animals in the RR-1 and RR-3 experiments were euthanized in space, this work suggested that space alone was the most likely cause for similar changes previously observed in liver samples from mice flown during the STS-135 experiments where animals returned live to Earth (Jonscher et al., 2016). The lipotoxic effect is stronger with duration and may have ramifications for astronauts’ health during long missions. This analysis did not include two flight and two ground animals from RR-1 as these animals were dissected immediately after euthanasia on-orbit as opposed to the rest which were first returned to Earth as intact frozen carcasses for later dissection.

Here we now compare RNA-sequencing (RNAseq) datasets generated from livers preserved using these distinct protocols. We find large changes in the transcriptome of tissues dissected and preserved on-orbit compared to tissues from mice euthanized on-orbit, preserved intact by freezing on-orbit, and dissected after return to Earth. To identify and mediate how the preservation method could have such a large effect on differential gene expression (DGE) results, we performed follow-on experiments to interrogate the role of euthanasia methods, tissue and carcass preservation protocols, and library preparation methods on DGE changes. Our findings have important implications for interpreting existing datasets and the design of future experiments.

## Results

### Preservation method is the primary driver of gene expression variance in RR-1 liver samples

To assess gene expression differences in liver samples from the RR-1 NASA Validation mission (Cadena et al., 2019; Globus and Galazka, 2015; Ronca et al., 2019), RNA was extracted from livers dissected from spaceflight (FLT) and ground control (GC) animals either immediately after euthanasia (immediate preservation, I), or from frozen carcasses after partial thawing (carcass preservation, C), and sequenced following polyA selection. Principal component analysis (PCA) revealed preservation method (C vs. I) as the primary driver of variance among samples rather than spaceflight (FLT vs. GC) (Figure 1A). Furthermore, there was an order of magnitude difference in the number of differentially expressed genes (DEGs) identified in FLT versus GC carcass samples than was observed in FLT versus GC immediate samples, and only 4 DEGs overlapped between the two preservation methods (Figure 1B). Gene set enrichment analysis of FLT versus GC C-(Figure 2A) and I-derived (Figure 2B) samples showed no overlap in enriched gene ontology (GO) terms (Figure 2C), showing that any gene expression changes in the liver as a result of spaceflight exposure were confounded by the sample preservation method used.

**Figure 1.**
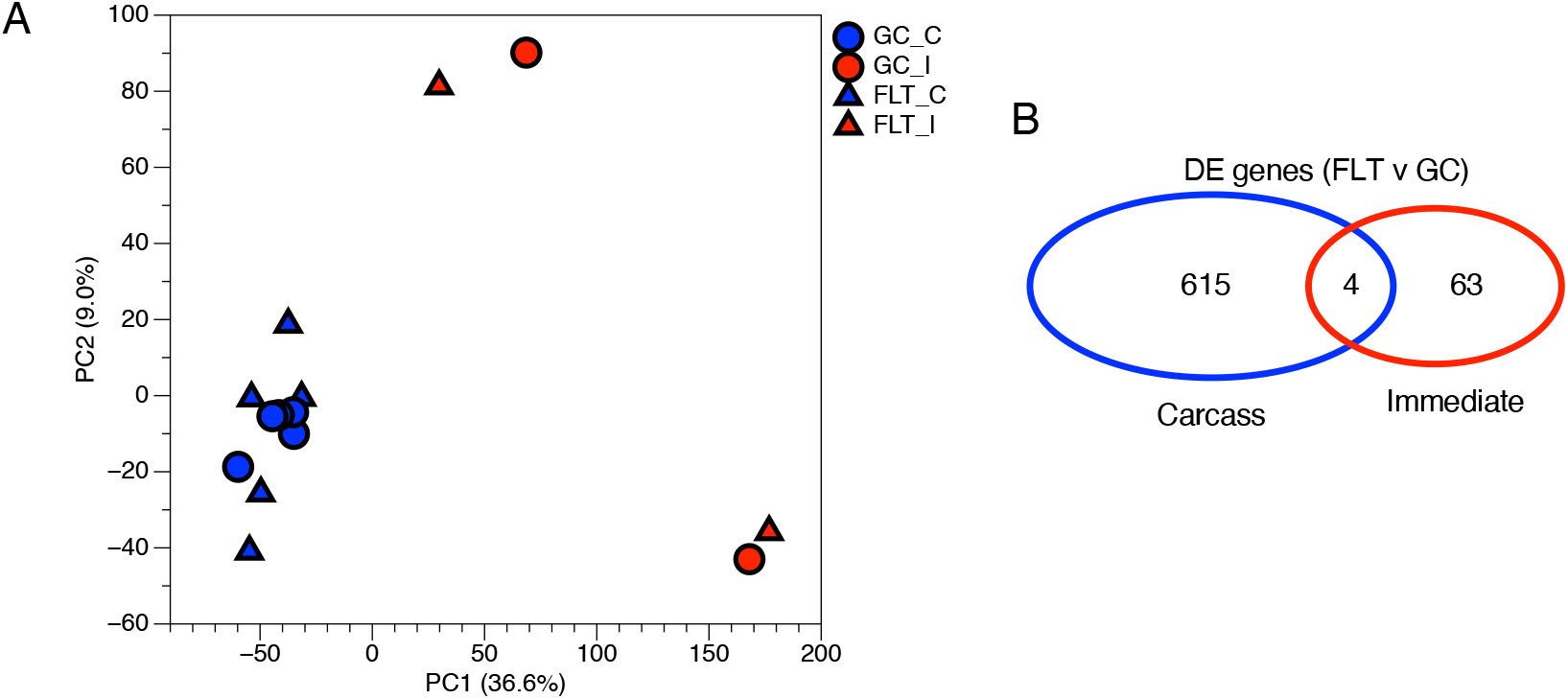
Gene expression differences in RR-1 NASA Validation flight liver samples prepared via polyA selection (GLDS-48). A) Principal component analysis of global gene expression in RR-1 NASA spaceflight (FLT) and respective ground control (GC) liver samples dissected immediately after euthanasia (I) or from frozen carcasses (C). Percent variance for each principal component (PC) is shown. B) Venn diagram showing the number of similar and unique differentially expressed genes, spaceflight (FLT) vs. ground control (GC), in Carcass (blue) and Immediate (red) samples (adj. p-value < 0.05).

**Figure 2.**
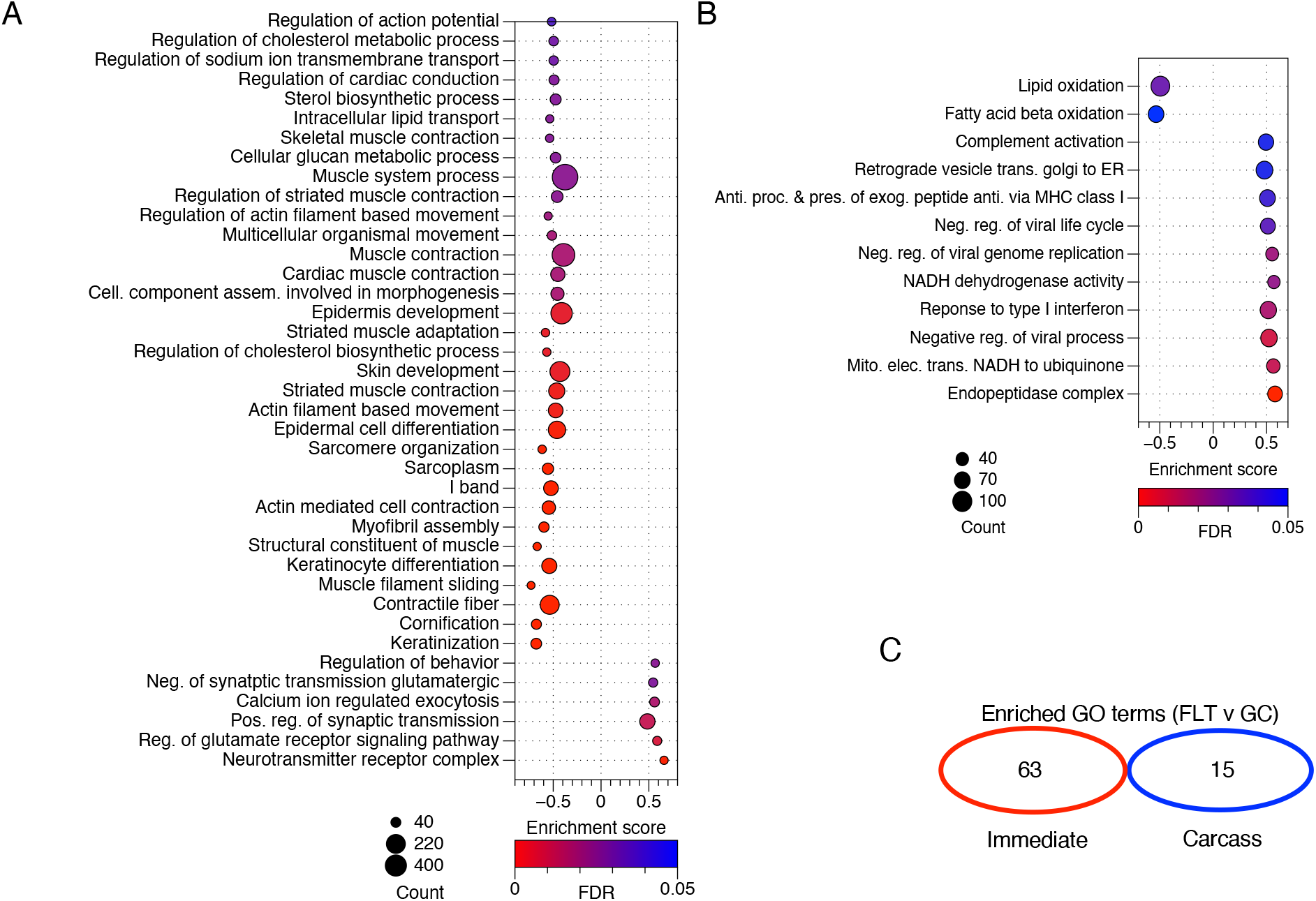
Enriched Gene Ontology (GO) terms between the RR-1 flight and ground groups. A) Enriched GO terms between the flight and ground control immediate samples (FLT-I vs. GC-I) identified by Gene Set Enrichment Analysis (get set permutation). B) Enriched GO terms between the flight and ground control carcass samples (FLT-C vs. GC-C) identified by Gene Set Enrichment Analysis (gene set permutation). In both A and B, positive or negative enrichment scores indicate higher expression in FLT-C or GC-C samples, respectively. Dot size indicates number of genes within GO term. Dot color indicates FDR. GO terms displayed met the thresholds of FDR < 0.05, NOM p < 0.01, gene set size > 40. C) Venn diagram of the number of enriched GO terms identified in Carcass (blue) and Immediate (red) samples when comparing FLT and GC samples. GO terms in Venn diagram met the threshold of FDR < 0.05, NOM p < 0.01.

Since livers from only 2 FLT and 2 GC animals in the RR-1 NASA Validation mission were preserved via the immediate method, RNA from livers prepared via the immediate method from two additional studies, the RR-1 CASIS mission (Cadena et al., 2019; Globus et al., 2015; Ronca et al., 2019) and a ground-based preservation and storage study (Choi et al., 2016; GeneLab, 2016) were also sequenced following polyA selection. Despite multiple different experimental factors in RR-1 NASA, RR-1 CASIS, and the ground-based preservation studies, PCA continued to show preservation method as the primary driver of variance among samples in these datasets (Figure S1).

### Carcass-preserved samples exhibit less uniform transcript coverage than immediate-preserved samples

To further investigate the observed differences in preservation method, RR-1 NASA FLT and GC liver samples derived from the carcass preservation method were grouped together (all-C) and FLT and GC liver samples derived from the immediate preservation method were grouped together (all-I). DGE was evaluated in all-C vs. all-I samples. Many more genes were differentially expressed in all-C vs. all-I samples than in either FLT-C vs. GC-C or in FLT-I vs. GC-I samples, further supporting preservation method as the primary driver of variance in RR-1 NASA liver samples (Figure 3A). Gene set enrichment analysis revealed that several of the gene ontologies enriched in carcass samples (when compared with immediate samples) involved RNA regulation and processing (Figure 3B). Despite similarly high RNA Integrity Number (RIN) values (Figure S2), carcass samples exhibited significantly less 5’ gene body coverage than immediate samples (Figure 3C&D).

**Figure 3.**
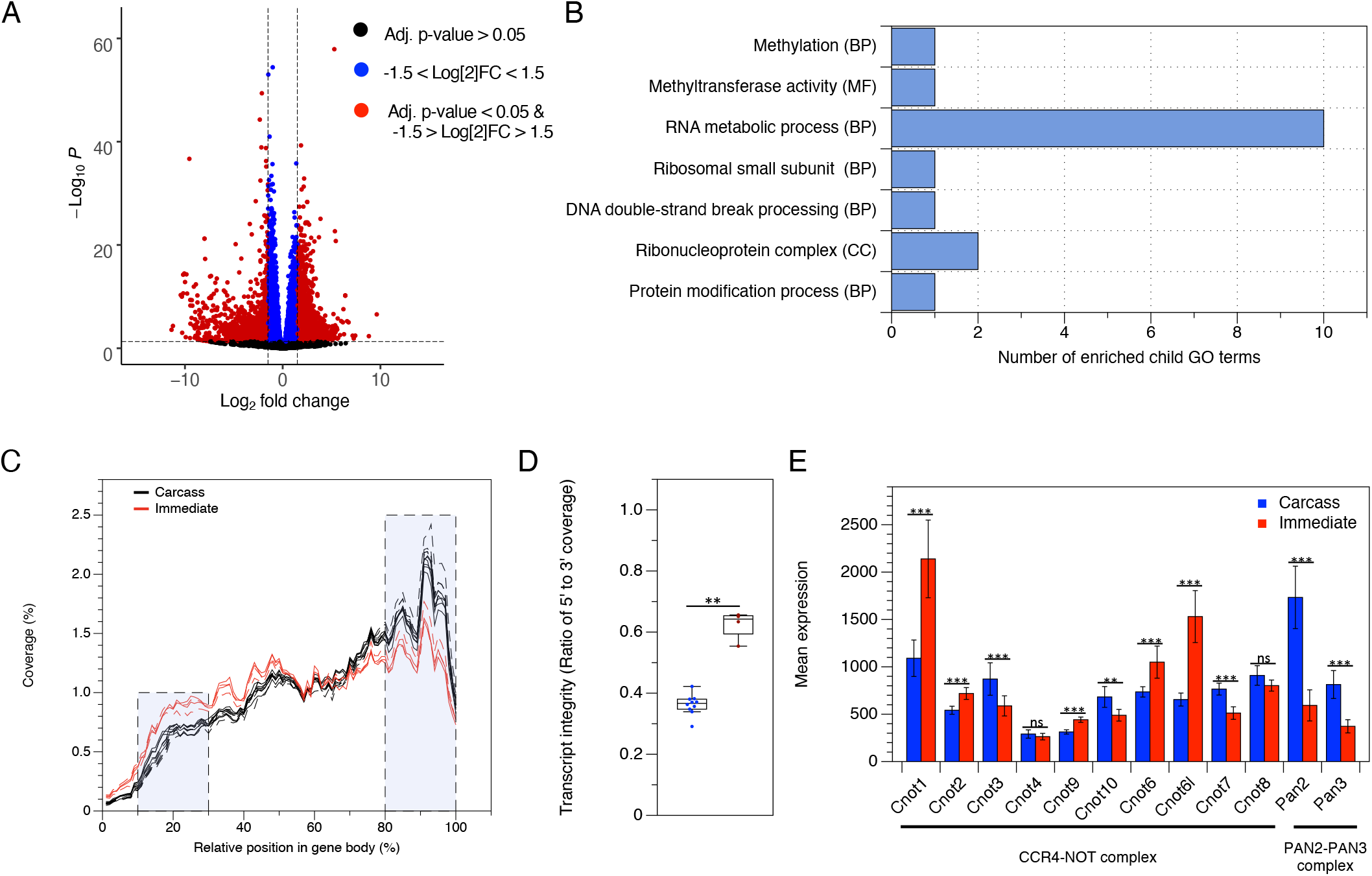
Gene expression changes and transcript integrity in Carcass vs. Immediate RR-1 NASA liver samples. A) Volcano plot showing 2,934 differentially expressed genes in all carcass (FLT-C and GC-C, all-C) versus all immediate (FLT-I and GC-I, all-I) samples (adj. p-value < 0.05 and 1.5 < Log2 fold change < −1.5). B) Common parent terms of enriched GO terms identified by Gene Set Enrichment Analysis of all carcass versus all immediate samples (phenotype normalized, FDR < 0.3, NOM p < 0.01). C) Gene body coverage in Carcass (black) and Immediate (red) FLT and GC samples. D) The percent coverage of the 5’ and 3’ shaded regions in panel C were used to calculate the 5’ to 3’ transcript integrity ratio for each sample. All Carcass and Immediate samples are grouped together (** = p < 0.01, Mann–Whitney U test). E) Average expression of poly(A) removal genes in Carcass (blue) and Immediate (red) groups from RNAseq data. Cnot1, Cnot2, Cnot3, Cnot4, Cnot9, Cnot10, Cnot6, Cnot6l, Cnot7, and Cnot8 are part of the CCR4-NOT complex. Pan2 and Pan3 are part of the PAN2-PAN3 complex. Error bars indicate standard deviation (* = adj. p < 0.05, ** = adj. p < 0.01, *** = adj. p < 0.001, ns = not significant, Wald test).

### Expression of genes involved in 5’-methylguanosine decapping and polyA removal is affected by preservation condition

Given the differences in gene body coverage between carcass and immediate samples, we evaluated the expression of 5’-methylguanosine decapping and polyA removal genes in these groups from the RNAseq data. In mammals, eight genes have decapping activity in vitro and/or in cells: Dcp2 (Nudt20), Nudt3, Nudt16, Nudt2, Nudt12, Nudt15, Nudt17, Nudt19. In addition, Dxo acts on partially capped mRNA’s (Grudzien-Nogalska and Kiledjian, 2017). Two of these genes — Dxo and Nudt2 — were significantly more expressed in the carcass samples, while another two — Nudt15 and Dcp2 (Nudt20) — were significantly more expressed in immediate samples (Figure S3). Removal of polyA tails from mRNA is catalyzed by two complexes. The first, CCR4-NOT, consists of CNOT1, CNOT2, CNOT3, CNOT4, CNOT9, CNOT10, CNOT6, CNOT6L, CNOT7, and CNOT8. The second, PAN2-PAN3, consists of PAN2 and PAN3 (Siwaszek et al., 2014). In the case of the 10 subunit CCR4-NOT complex, we observed 5 genes that were more highly expressed in the immediate group (Cnot1, Cnot2, Cnot9, Cnot6, Cnot6l) and 3 that were more highly expressed in the carcass group (Cnot3, Cnot10, Cnot7) (Figure 3E). In the case of the PAN2-PAN3 complex, both Pan2 and Pan3 were more highly expressed in the carcass group (Figure 3E).

### Samples sequenced following ribodepletion exhibit more uniform transcript coverage than samples prepared with polyA selection

The polyA selection library preparation method, which was initially used to evaluate gene expression differences in RR-1 NASA Validation mission liver samples requires intact RNA to minimize bias (Kumar et al., 2017; Petrova et al., 2017). Since our data suggest that the carcass samples were more degraded than the immediate samples (Figure 3C&D), the total RNA isolated from the RR-1 carcass liver samples was used to prepare libraries with the ribodepletion method to minimize transcript integrity bias, then re-sequenced. PCA showed a more distinct separation of FLT and GC carcass samples when the samples were prepared via the ribodepletion method (Figure 4A) than by polyA selection (FLT-C and GC-C samples in Figure 1A). DEGs were identified in FLT vs. GC carcass samples prepared with the ribodepletion method and compared with those from poly-A prepared carcass samples. Although hundreds of DEGs in FLT vs. GC carcass samples overlap between ribodepleted and polyA prepared samples, more DEGs were identified in FLT vs. GC samples prepared with the ribodepletion method (Figure 4B), suggesting this method may be more sensitive. There was no overlap of enriched gene ontology terms in FLT vs. GC samples processed by ribodepletion and polyA enrichment (Figure 4C&D).

**Figure 4.**
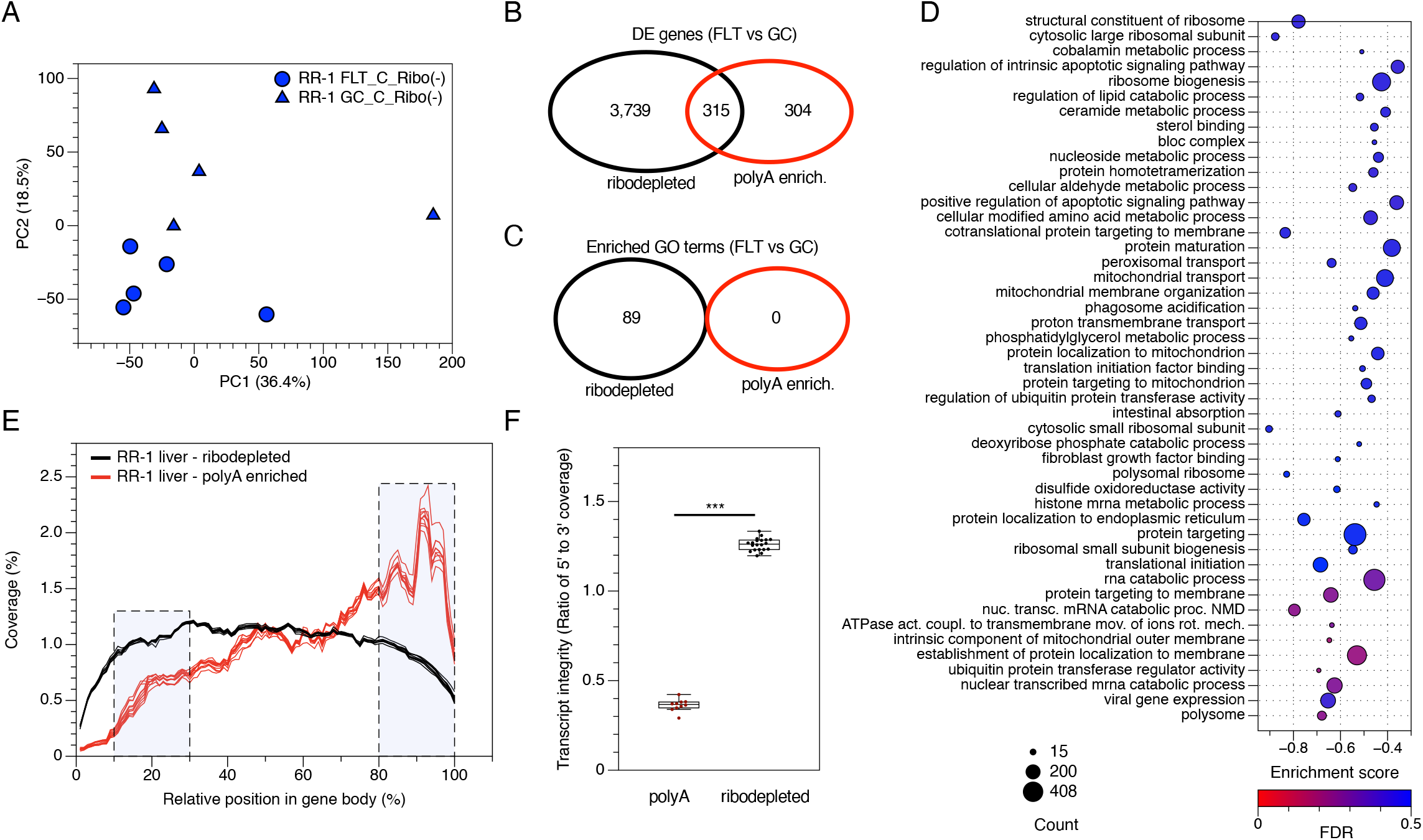
Evaluation of gene expression and transcript integrity in RR-1 NASA carcass-dissected FLT and GC samples prepared via polyA selection and ribodepletion methods. A) Principal component analysis of global gene expression in RR-1 NASA spaceflight (FLT) and ground control (GC) liver samples dissected from frozen carcasses and prepared via ribodepletion. Percent variance for each principal component (PC) is shown. B) Venn diagram of differentially expressed (DE) genes between spaceflight (FLT) and ground control (GC) samples prepared with ribodepletion (black) or polyA selection (red) methods (adj. p-value < 0.05). C) Venn diagram of the number of similar and unique enriched GO terms identified in ribodepleted (black) and polyA selected (red) prepared samples (NOM p < 0.01, FDR < 0.5, phenotype permutation). D) Enriched GO terms between the flight and ground control carcass samples prepared with ribodepletion identified by Gene Set Enrichment Analysis (phenotype permutation). Positive or negative enrichment scores indicate higher expression in FLT or GC samples, respectively. Dot size indicates number of genes within GO term. Dot color indicates FDR. GO terms displayed met the thresholds of FDR < 0.5, NOM p < 0.01, 1.6 < NES < −1.6. E) Gene body coverage of ribodepleted and polyA-selected FLT and GC carcass samples. F) The percent coverage of the 5’ and 3’ shaded regions were used to calculate the transcript integrity ratio for each sample (*** = p < 0.001, Mann–Whitney U test).

Next, transcript integrity was evaluated in the ribodepletion-prepared FLT and GC carcass samples and compared with polyA selection-prepared carcass samples. Samples prepared with the polyA selection method exhibited less coverage of the 5’ portion of transcripts compared to ribodepletion-prepared samples (Figure 4E&F). Thus ribodepletion was used to further investigate the effects of preservation method (Carcass vs. Immediate) on gene-expression in a ground study.

### Total RNA sequencing mitigates the impact of preservation method on gene expression changes in the liver

We designed a ground-based tissue preservation study to determine the best approach to mitigate the impact of preservation method on gene expression, and to identify other confounding variables important for interpreting data from other RR missions. We tested euthanasia and preservation techniques used in different RR missions and compared them to standard laboratory protocols for tissue preservation. In addition to liver samples, we also analyzed quadriceps to determine whether sample preservation methods also confounded DGE analysis in this tissue.

Mice of the same age, sex, strain, and source as those used in the RR-1 NASA Validation mission were used in the ground-based tissue preservation study. Mice were evenly divided into one of 6 groups as shown in Figure S4. The mice in groups 1-4 were euthanized with pentobarbital/phenytoin (Euthasol^®^) as in RR-1 (Choi et al., 2020), then subjected to various preservation protocols to evaluate the phenomena observed in RR-1 NASA carcass and immediate liver samples. Livers and quadriceps were dissected immediately after euthanasia from mice in group 1. These tissues were divided into thirds and preserved in one of three ways: 1) freezing in dry ice to mimic the cold block that was used to freeze the immediate liver samples in the RR-1 NASA mission, 2) submersion in LN2, or 3) with RNA*later*™. After preservation, all tissues were stored at − 80 °C until further processing.

Although it is common practice to dissect mice immediately after euthanasia, due to limitations in crew time for spaceflight experiments, immediate dissection is not always possible. Thus, most tissues are preserved *in situ* within the carcass. We therefore sought to determine the most effective way to preserve carcasses that would minimize unintended gene expression changes in tissues preserved *in situ*. Mice in groups 2-4 were used to test three different carcass preservation methods: 1) slow freezing in dry ice (DI) to mimic the most common method of carcass preservation used in RR missions to date, 2) snap freezing by submersion in liquid nitrogen (LN2), and 3) segmenting the carcass into thirds and preserving in RNA*later*™, mimicking the preservation method used in the Rodent Research-7 mission. After preservation, all carcasses were stored at −80 °C until further processing.

Carcasses from mice in groups 2-4 were partially thawed, and quadriceps and livers were dissected, then snap frozen, and stored at −80 °C until RNA extraction to mimic the protocol most commonly implemented when carcasses return from spaceflight missions, including RR-1. A summary of all liver and quadriceps tissues evaluated in the present ground-based tissue preservation study are summarized in Tables S1 and S2, respectively. Total RNA was extracted from all liver and quadriceps tissue samples and prepared for sequencing using the ribodepletion method and sequenced.

Global gene expression and transcript integrity were evaluated in liver samples from groups 1-4 to identify differences in DGE resulting from the carcass and immediate preservation protocols. PCA showed overlap among immediate samples despite differences in tissue preservation methods (Figure 5A). Similar to RR-1 carcass and immediate samples (Figures 1A & S1), the dry ice-preserved carcass and immediate samples, which mimic the RR-1 preservation conditions, clustered away from each other, albeit to a much lesser degree than that observed with the RR-1 samples (Figure 5A). Furthermore, the dry ice-preserved carcass samples exhibited less 5’ gene body coverage than the dry ice-preserved immediate samples (Figure 5B). Although this observation is consistent with that observed in RR-1 carcass and immediate samples (prepared using the polyA selection method) (Figure 3C&D), the difference was less dramatic. Therefore, using the ribodepletion method appears to partially alleviate the differences observed in transcript integrity between carcass and immediate samples.

**Figure 5.**
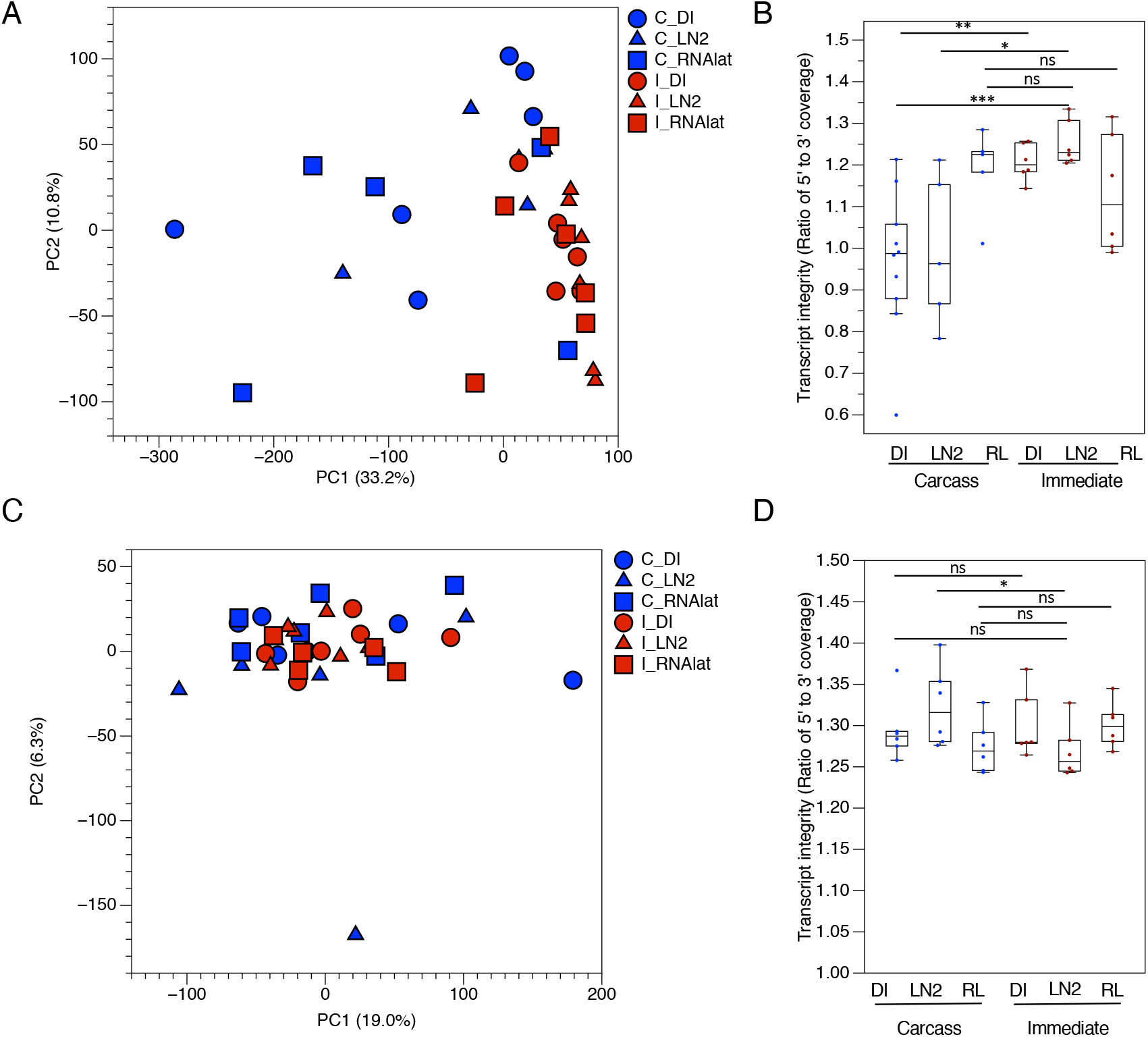
Gene expression and transcript integrity analysis of preservation methods for liver and quadriceps samples. Liver and quadriceps samples were dissected from mice immediately after euthanasia with pentobarbital/phenytoin (Euthasol^®^) then preserved in dry ice (I_DI), liquid nitrogen (I_LN2), or RNA*later*™ (I_RL) before RNAseq analysis. Alternatively, liver and quadriceps samples were dissected from partially thawed frozen carcasses of mice that were euthanized with pentobarbital/phenytoin (Euthasol^®^) then preserved in dry ice (C_DI), liquid nitrogen (C_LN2), or segmented into thirds and preserved in RNA*later™* (C_RL) before RNAseq analysis. A) Principal component analysis of liver samples. Percent variance for each principal component (PC) is shown. B) Uniformity of gene body coverage in liver samples. C) Principal component analysis of quadriceps samples. Percent variance for each principal component (PC) is shown. D) Uniformity of gene body coverage in quadriceps samples. (*** = p < 0.001, ** = p < 0.01, * = p < 0.05, ns = no significance, Mann–Whitney U test).

### Carcass preservation by LN2 or RNAlater™ immersion most closely mimic standard tissue preservation protocols

Livers dissected from carcasses preserved in either RNA*later*™ or LN2 exhibit more overlap with immediate preserved liver samples than those from carcasses preserved in dry ice (Figure 5A). Unlike livers dissected from slow (dry ice) or snap (LN2) frozen carcasses, livers dissected from carcasses preserved in RNA*later*™ showed no difference in 5’ to 3’ transcript coverage when compared with livers dissected immediately after euthanasia (Figure 5B). These data suggest that carcass segmentation and preservation in RNA*later*™ may protect the liver from transcript degradation when preserved *in situ*.

We next assessed the effects of various carcass freezing methods on gene expression changes in the liver when compared with livers that were dissected immediately after euthanasia then preserved in either RNA*later*™ or LN2. Only a few genes were differentially expressed between livers dissected immediately and preserved on dry ice, in RNA*later*™, or in LN2 (Table S3). Similarly, pairwise gene set enrichment analysis showed no significantly enriched GO terms between these tissue preservation methods (Table S3), suggesting that for immediately dissected livers, the tissue preservation method had minimal impact on gene expression. Livers dissected from slow (dry ice) frozen carcasses, which most closely mimics the carcass preservation method used in RR-1 NASA and several other RR missions (including RR-3 and Rodent Research-6), exhibited the most DEGs when compared with immediately dissected livers preserved in either LN2 or RNA*later*™ (Tables 1 and S4, respectively). In contrast, livers dissected from carcasses preserved in either RNA*later*™ or LN2 exhibited hundreds fewer DEGs when compared with immediately dissected livers preserved in either LN2 or RNA*later*™ (Tables 1 and S4, respectively). These data indicate that carcass segmentation and preservation in RNA*later*™ or preserving carcasses by submersion in LN2 more closely mimic the common preservation methods used in terrestrial laboratories, than does the slow freeze carcass preservation method used in the RR-1 NASA Validation study.

**Table 1.** Comparisons of carcass preservation methods to immediate liquid nitrogen method on gene expression in livers. The number of differentially expressed genes (DEG) and enriched gene ontology (GO) terms identified by Gene Set Enrichment Analysis (phenotype permutation) were evaluated pairwise in liver samples from different carcass preservation methods compared with immediate samples preserved in liquid nitrogen. For GO terms, number on the left corresponds to the group to the left of the ‘vs.’, and number on the right corresponds to the group to the right of the ‘vs.’ in the “Comparison” column. n numbers, p values, log2 fold changes, and FDR values are indicated. Euth=euthanasia by pentobarbital/phenytoin, I=tissue dissected immediately after euthanasia, C=tissue dissected from frozen carcass that has been partially thawed, DI=dry ice, LN2=liquid nitrogen, RL= RNA*later*™. Data are from GLDS-235.

| Comparison | # DEG<br>(adj. p < 0.05) | # DEG<br>(adj. p < 0.05 &<br>\|Log2 FCI > 1.5) | # Enriched GO<br>terms<br>(NOM p < 0.01) | # Enriched GO<br>terms<br>(FDR < 0.5 &<br>NOM p < 0.01) | # Enriched GO<br>terms<br>(FDR < 0.25 &<br>NOM p < 0.01) |
| --- | --- | --- | --- | --- | --- |
| Euth_C_DI<br>(n=6) vs.<br>Euth_I_LN2<br>(n=6) | 3798 | 3143 | 16, 0 | 0, 0 | 0, 0 |
| Euth_C_LN2<br>(n=5) vs.<br>Euth_I_LN2<br>(n=6) | 784 | 515 | 30, 0 | 0, 0 | 0, 0 |
| Euth_C_RL<br>(n=5) vs.<br>Euth_I_LN2<br>(n=6) | 2118 | 1952 | 22, 0 | 0, 0 | 0, 0 |

### The impact of preservation method on gene expression is tissue dependent

To determine if the observed differences in gene expression due to carcass preservation method is unique to the liver, gene expression and transcript integrity was also evaluated in quadriceps from mice in groups 1-4 (Figure S4 and Table S2). PCA showed more overlap among carcass and immediate quadriceps samples (Figure 5C) than among carcass and immediate liver samples (Figure 5A), suggesting that gene expression in the quadriceps is less sensitive to preservation methods. Unlike liver samples, almost no significant differences were observed in 5’ to 3’ gene body coverage in quadriceps samples prepared using different preservation methods (Figure 5D).

Fewer DEGs were identified in carcass vs. immediate quadriceps samples than carcass vs. immediate liver samples for almost every preservation method tested (Tables 2, S5, S6), further supporting that gene expression in the quadriceps is less sensitive to different types of preservation methods. Although there are fewer differences over-all, similar to what was observed in liver samples, cutting the carcass into thirds then preserving in RNA*later™* resulted in the fewest DEGs when compared with immediate dissection followed by tissue preservation in LN2 or RNA*later™* (Tables 2 and S6).

**Table 2.**
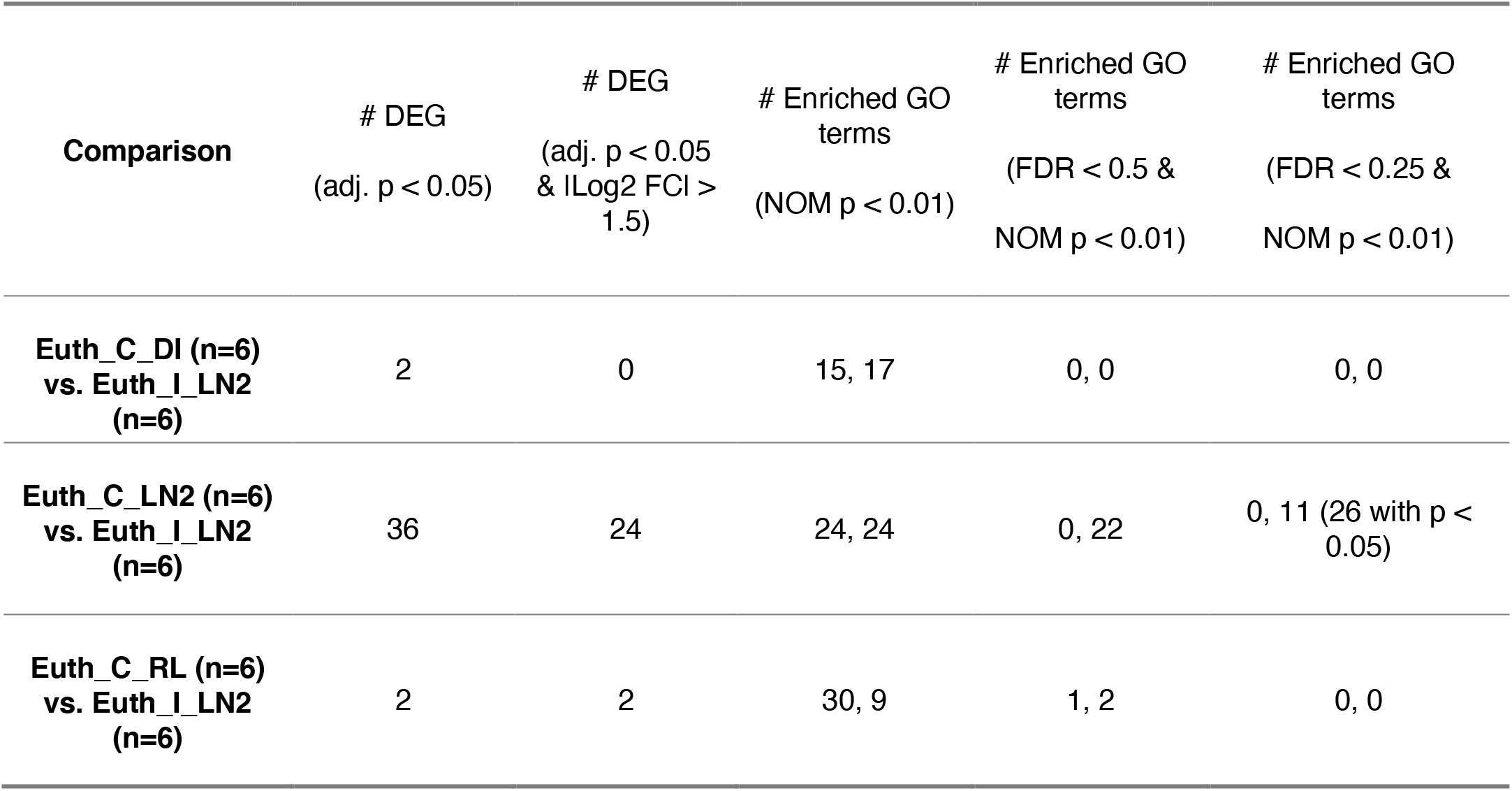
Comparisons of carcass preservation methods to immediate liquid nitrogen method on gene expression in quadriceps. The number of differentially expressed genes (DEG) and enriched gene ontology (GO) terms identified by Gene Set Enrichment Analysis (phenotype permutation) were evaluated pairwise in quadriceps samples from different carcass preservation methods compared with immediate samples preserved in liquid nitrogen. For GO terms, the first number corresponds to the group to the left of the ‘vs.’, and the second number corresponds to the group to the right of the ‘vs.’ in the “Comparison” column. n numbers, p values, log2 fold changes, and FDR values are indicated. Euth=euthanasia by pentobarbital/phenytoin, I=tissue dissected immediately after euthanasia, C=tissue dissected from frozen carcass that has been partially thawed, DI=dry ice, LN2=liquid nitrogen, RL=RNA*later*™. Data are from GLDS-236.

| Comparison | # DEG<br>(adj. p < 0.05) | # DEG<br>(adj. p < 0.05<br>& \|Log2 FCI > 1.5) | # Enriched GO<br>terms<br>(NOM p < 0.01) | # Enriched GO<br>terms<br>(FDR < 0.5 &<br>NOM p < 0.01) | # Enriched GO<br>terms<br>(FDR < 0.25 &<br>NOM p < 0.01) |
| --- | --- | --- | --- | --- | --- |
| Euth_C_DI (n=6)<br>vs. Euth_I_LN2<br>(n=6) | 2 | 0 | 15, 17 | 0, 0 | 0, 0 |
| Euth_C_LN2 (n=6)<br>vs. Euth_I_LN2<br>(n=6) | 36 | 24 | 24, 24 | 0, 22 | 0, 11 (26 with p < 0.05) |
| Euth_C_RL (n=6)<br>vs. Euth_I_LN2<br>(n=6) | 2 | 2 | 30, 9 | 1, 2 | 0, 0 |

### Gene expression in select tissues was not affected by the method of euthanasia

Since the most common euthanasia method used in RR missions to date is intraperitoneal (IP) injection of ketamine/xylazine and the most common euthanasia method used in standard laboratories is CO_2_ inhalation, these methods were used to euthanize mice in groups 5 and 6, respectively, to determine if euthanasia method is another confounding variable that could affect gene expression in select tissues (Figure S4, Tables S1 and S2). Gene expression was evaluated in livers and quadriceps dissected from mice in groups 2, 5, and 6 (Figure S5A-D). PCA showed no distinct differences in global gene expression in liver (Figure S5A) or quadriceps (Figure S5B) samples dissected from mice euthanized with different methods. Pairwise differential gene expression analysis and gene set enrichment analysis also identified few, if any, DEGs and enriched GO terms among liver (Figure S5C) and quadriceps (Figure S5D) samples. These data suggest that the types of euthanasia methods evaluated here do not impact gene expression in select tissues.

## Discussion

Herein, we show that protocols used to preserve mouse carcasses on-orbit have large effects on gene expression patterns as measured by RNAseq. Indeed, changes in gene expression due to preservation condition overwhelmed those due to spaceflight. Gene set enrichment analysis showed that many GO terms enriched due to carcass preservation were involved in RNA processing. This correlated with reduced transcript integrity (relatively poor coverage of the 5’ end of transcripts) in samples from carcasses preserved on-orbit when these were sequenced with a polyA enrichment RNAseq protocol.

While RNAseq following polyA selection can more efficiently quantify gene expression (Kumar et al., 2017), ribodepletion methods are more effective on degraded RNA samples (Li et al., 2014; Schuierer et al., 2017). However, while the RNA used in this study was of good quality (RIN > 7) we observed a severe bias in transcript coverage following polyA selection depending upon the tissue preservation condition utilized. Specifically, samples taken from carcasses that were slow-frozen on-orbit exhibited a lower 5’ to 3’ coverage ratio as compared to samples taken from immediately dissected tissues. While resequencing of the carcass flight samples with a ribodepletion protocol produced a more even 5’ to 3’ coverage ratio, our follow-on studies that directly compared slow carcass freezing to immediate dissection revealed a similar (albeit reduced) 5’ to 3’ coverage bias. Taken together, this suggests that slow carcass freezing causes transcript degradation that in-turn leads to reduced 5’ coverage.

mRNA degradation starts with the removal of the polyA tail, at which point degradation continues either from the 3’ end *via* the exosome complex, or the 5’ end following removal of the 5’-methylguanosine cap. Deadenylation of cytoplasmic mRNA is the rate limiting step in mRNA degradation and is catalyzed by one of two complexes. The CCR4-NOT complex, which consists of 10 subunits (CNOT1, CNOT2, CNOT3, CNOT4, CNOT6, CNOT6l, CNOT7, CNOT8, CNOT9, CNOT10), and the PAN2-PAN3 deadenylation complex consisting of two subunits (PAN2, PAN3) (Siwaszek et al., 2014). We observed transcriptional changes to multiple subunits in each of these complexes when comparing carcass and immediate samples. Most striking was the coordinate upregulation of both Pan2 and Pan3 in the carcass samples from RR-1, which suggest an increase in PAN2-PAN3 deadenylation activity, which could result in loss of polyA tails in some transcripts. This could lead to poor mRNA capture by our polyA enrichment protocol and result in some of the differences seen been the polyA-enrichment and ribodepletion protocols.

Three proteins - Dcp2 (Nudt20), Nudt3, Nudt16 - have decapping activity both in vitro and in cells, while an additional five – Nudt2, Nudt12, Nudt15, Nudt17, Nudt19 – have decapping activity in vitro. In addition, the Dxo family of proteins acts on partially capped mRNA’s (Grudzien-Nogalska and Kiledjian, 2017). While regulation of these proteins is complex and involves subcellular localization and post-translation modification, we observed evidence for altered expression of these decapping enzymes: Dxo and Nudt2 were more abundant in carcass samples, while Nudt15 and Dcp2 (Nudt20) were more abundant in immediate samples. While these changes are not coordinated, they do point to altered decapping activity within the carcass samples. As decapping proceeds mRNA degradation via the 5’ exonuclease, XRN1, this could alternatively explain the relatively poor 5’ transcript coverage seen in both polyA enriched and ribodepleted carcass liver samples. Additional experimentation will be necessary to confirm the changes to decapping and deadenylation enzymes seen here and to understand their role in the 5’ to 3’ coverage bias observed.

We observed a marked difference in the 5’ to 3’ coverage bias between liver and quadriceps samples. Whereas liver samples were sensitive to carcass preservation *via* slow- or snap-freeze, quadricep samples were not. There are a number of possible explanations for this. First, it could be due to the surface exposure of the quadriceps, which would lead to more rapid quenching of biochemical processes. Second, inherent differences in the transcript pool, mRNA half-lives, and enzymatic complement of liver and quadriceps could offer a biological answer. While we cannot distinguish between these mechanisms, our observations are consistent with previous results showing that post-mortem changes to mRNA is tissue-dependent (Inoue et al., 2002; Lee et al., 2005; Miyatake et al., 2004).

The poor transcript integrity in slow frozen carcasses sequenced with a polyA enrichment protocol was not evident in pre-sequencing QC analyses. Indeed, all samples had RIN values > 7 and there was no correlation between the gene expression differences and RIN. This distinguishes our results from previous studies showing a strong correlation between RIN values and loss of 5’ coverage (Davila et al., 2016; Sigurgeirsson et al., 2014). Therefore, additional pre-sequencing QC analyses capable of detecting these issues would be useful. Low throughput sequencing is rapid, decreasing in cost, and being adopted as a QC step, but does not provide the coverage necessary to detect the biases seen here.

Alternatively, if effective pre-sequencing QC metrics cannot be developed, a number of analytical approaches could be utilized. In one category are methods that calculate additional metrics such as mRIN (Feng et al., 2015) and TIN (Wang et al., 2016) to allow assessment transcript integrity and exclusion of problematic samples. In a second category are processes that account for variable transcript integrity by considering only reads that occur near the 3’ end of transcripts (Sigurgeirsson et al., 2014), controlling for the effects of RIN using a linear model framework (Gallego Romero et al., 2014), or by calculating idealized coverage curves on a gene-by-gene basis and using these for normalization (Xiong et al., 2019). Additional analyses are necessary to determine if these approaches can mitigate the issue observed here.

While we do not have a complete picture of the mechanisms resulting in the apparent gene expression change resulting from slow carcass freezing, we were able to identify effective mitigation strategies. Foremost among these is the utilization of a ribodepletion protocol in place of polyA enrichment. In this study, ribodepletion resulted in more even gene body coverage and was not as sensitive to slow freezing of carcasses. This is in agreement with previous studies which found that ribodepletion is less prone to bias introduced by poor RNA quality (Li et al., 2014) and less prone to 3’ coverage bias (Schuierer et al., 2017). Beyond this, we found that two carcass preservation methods generated acceptable results, with few DEGs and enriched GO terms when compared to the immediate dissection of tissues and preservation in liquid nitrogen — the *de facto* gold standard. The first is rapid freeze of carcasses in liquid nitrogen and subsequent storage at −80 °C, followed by partial thaw, dissection and tissue preservation in liquid nitrogen. While this led to some loss of 5’ transcript coverage, it had the fewest DEGs (515, adj. p < 0.05 & |Log2 FC| > 1.5) and no enriched GO terms (FDR < 0.25, NOM < 0.01) when compared to immediate dissection. Alternatively, segmentation of carcasses and immersion in RNA*later* and subsequent storage at −80 °C, followed by partial thaw, dissection and tissue preservation in liquid nitrogen resulted in better maintenance of 5’ transcript coverage but an increased number of DEGs (1952, adj. p < 0.05 & |Log2 FC| > 1.5) although no GO terms were enriched (FDR < 0.25, NOM < 0.01). As euthanasia protocols can change serum biomarkers (Pierozan et al., 2017) and mRNA expression levels (Staib-Lasarzik et al., 2014), we were reassured to find that the euthanasia protocols used here did not affect gene expression in liver or quadriceps.

To conclude, our results indicate that care must be taken in choosing sample preservation protocols that preserve transcriptional patterns and other embedded information, but that are also feasible in resource constrained environments such as those found in space.

## STAR Methods

### Rodent Research-1 (RR-1) Study

#### Spaceflight Mission

Rodent Research-1 (RR-1) was the first mission in which animals were maintained on the ISS for a long duration mission in the Rodent Habitat modified from heritage Animal Enclosure Module (AEM) hardware. Complete details were published previously (Choi et al., 2020). In short, RR-1 consisted of two experiments: ISS National Lab study (RR-1 CASIS) and NASA Validation study (RR-1 NASA). In the ISS National Lab Study, ten 32-week-old female C57BL/6NTac mice (Taconic Biosciences, Rensselaer, NY) were flown to space for 20-21 days then euthanized via IP injection of pentobarbital/phenytoin (Euthasol^®^) and dissected onboard the ISS. Livers were dissected then inserted into cryovials, which were then frozen in a cold stowage container that was pre-chilled to − 130 °C before transferring to the Minus Eighty-Degree Laboratory Freezer (MELFI) at the end of each dissection session (-80 °C). In the NASA Validation study, ten 16-week-old female C57BL/6J mice (Jackson laboratories, Bar harbor, ME) were flown to the ISS for 37 days before euthanasia and subsequent dissection. Due to crew time constraint, only two (out of ten) mice were dissected immediately after euthanasia via IP injection of pentobarbital/phenytoin (Euthasol^®^) to recover spleen and liver tissues on the ISS. Isolated livers were preserved by using the same method as the ISS National Lab study livers. The remaining eight animals were euthanized, then intact carcasses were wrapped in aluminum foil, put in Ziploc bags, placed in a pre-chilled cold stowage container and stored in the MELFI. For both the ISS National Lab study and the NASA Validation study, there were respective cohorts of age-matched basal animals which were euthanized one day after launch as a baseline control as well as age-matched ground control animals kept in an ISS Environmental Simulator at Kennedy Space Center (KSC) on a 4-day delay to mimic spaceflight conditions. In addition, the NASA Validation study also had a cohort of age-matched vivarium control animals that were housed in the vivarium cages and followed the same experimental timeline and process as the spaceflight animals. A timeline indicating major events in the RR-1 mission is shown in Supplemental Figure 6.

#### Sample Collection

The frozen intact carcasses from the NASA Validation study were partially thawed then dissected at NASA Ames Research Center upon return to Earth. One lobe of liver from each carcass was removed, immediately homogenized in RLT buffer (Qiagen, Valencia, CA) followed by snap freezing the tissue homogenates in LN2. Quadriceps were snap frozen upon collection. Tissues were stored at −80 °C until extraction.

#### RNA Isolation

RNA was isolated from all liver and quadriceps samples using the following methods. For the liver samples, RNA was extracted with the AllPrep DNA/RNA Mini Kit (Qiagen, Valencia, CA) following the manufacturer’s protocol. Briefly, homogenization buffer was made by adding 1:100 volume ratio of beta-mercaptoethanol to RLT buffer and kept on ice until use. Approximately 30 mg of tissue was cut using a sterile scalpel and immediately placed in 800 μL of the RLT buffer solution. Each sample was then homogenized for approximately 20 seconds at 21,000 RPM using a Polytron PT1300D handheld homogenizer with a 5 mm standard dispersing aggregate tip (Kinematica, Bohemia, NY). Homogenates were centrifuged for 3 minutes at room temperature at 15,000 RPM to remove cell debris. The supernatant from each sample was used to isolate and purify RNA following the manufacturer’s protocol including on-column DNase treatment with RNase-free DNase (Qiagen, Valencia, CA). RNA was eluted twice per sample in 30 μL RNase- and DNase-free H2O per elution. For quadriceps samples, RNA was extracted using TRIzol reagents (Thermo Fisher Scientific, Waltham, MA) according to the manufacturer’s protocol, and the isolated RNA samples were then treated on column with RNase-free DNase (Qiagen, Valencia, CA) and RNeasy Mini kit (Qiagen, Valencia, CA). Concentration and absorbance ratios of all the isolated liver and quadriceps RNA samples were measured using the NanoDrop 2000 spectrophotometer (Thermo Fisher Scientific, Waltham, MA). RNA quality was assessed using the Agilent 2100 Bioanalyzer with the Agilent RNA 6000 Nano Kit or Agilent RNA 6000 Pico Kit (Agilent Technologies, Santa Clara, CA).

#### Library Preparation and RNA-Sequencing

Samples with RNA Integrity Number (RIN) of 6 or above were sent to the University of California (UC), Davis Genome Center where the libraries were constructed and RNA- sequencing was performed. All the RR-1 RNA-sequencing data analyzed in this manuscript were obtained from the NASA GeneLab Data Repository (https://genelab.nasa.gov/), including GLDS-47, GLDS-48, and GLDS-168. The RR-1 liver RNA samples were sequenced twice. First, libraries were generated using the Illumina TruSeq Stranded RNA library prep kit (Illumina, San Diego, CA) after polyA selection, and sequencing was done with 50 bp single end reads on the Illumina HiSeq 3000 platform (GLDS-47 and GLDS-48). Second, selected RNA samples were spiked in with ERCC ExFold RNA Spike-In Mixes (Thermo Fisher Scientific, Waltham, MA) before shipping to the UC Davis Genome Center. Ribosomal RNA was removed with the Illumina RiboZero Gold ribodepletion kit then RNA sequencing libraries were constructed using the KAPA RNA HyperPrep kit (Roche, Basel, Switzerland) and the sequencing was done with 150 bp paired end reads on the Illumina HiSeq 4000 platform (GLDS-168).

### Ground-based Tissue Preservation Study

#### Animals

20- to 21-week-old female C57BL/6J mice (Jackson laboratories, Bar harbor, ME) were shipped to the NASA Ames Research Center Animal Care Facility and were randomly housed in the standard vivarium cages with up to five mice per cage. The animals were acclimated for five days before the start of procedures to ensure recovery from the transportation stress. During acclimation, the animals were maintained on a 12h light/dark cycle and were provided with standard chow and water access *ad libitum*. One day before euthanasia, animal body weights were measured and used to distribute the animals into six groups (n=6/group) with similar average body weights. Animal health status, water and food intake were monitored daily. The study followed recommendations in the Guide for the Care and Use of Laboratory Animals (2011) and was approved on February 8, 2018 by the Institutional Animal Care and Use Committee (IACUC) at NASA Ames Research Center (Protocol number NAS-17-006-Y1).

#### Animal Euthanasia and Dissection

The detailed descriptions and rationale of each group are as follows as well as outlined in Figure S4 and Tables S1 and S2. Group 1 (M1-M6) animals were euthanized by intraperitoneal injection of pentobarbital/phenytoin (Euthasol^®^) (80 mg in 0.2 ml) (Virbac, West Lake, TX) followed by cervical dislocation. Dissection was performed immediately after euthanasia without freezing the carcasses. Left lobes of livers and quadriceps were subdivided into three sections and each tissue section was preserved either by freezing on dry ice, snap freezing in liquid nitrogen, or preserved in RNA*later*™ solution (Thermo Fisher Scientific, Waltham, WA). For the tissue sections preserved in RNA*later*™, tissue sections were submerged in RNA*later*™ at 4 °C for 3 days then frozen and stored at −80 °C. Note that this is the only group of animals that were dissected upon euthanasia. Carcasses from animals in subsequent groups were preserved intact using various methods then dissected at a later date. Group 1 tissue sections that were preserved by freezing on dry ice most closely mimics the process that was used to generate RR-1 NASA and CASIS immediate samples.

Group 2 (M7-M12) animals were euthanized by intraperitoneal injection of pentobarbital/phenytoin (Euthasol^®^) followed by cervical dislocation. The carcasses were wrapped in foil and preserved by freezing on dry ice, similar to the intact carcass preservation method used for the RR-1 NASA Validation Study. Once frozen, the carcasses were stored at −80 °C. On the day of dissection, mouse carcasses were removed from the −80 °C freezer and thawed at room temperature for 15 to 20 minutes prior to dissection. Left lobes of livers were removed and divided into two: one piece was snap frozen in liquid nitrogen then stored at −80 °C; the other piece was homogenized in RLT buffer (Qiagen, Valencia, CA) and the tissue homogenate was snap frozen in liquid nitrogen then stored at −80 °C for 70 days before RNA extraction to simulate the process used to generate the RR-1 NASA “carcass” liver samples. This extended storage did not result in a substantial number of DEG (Figure S5E&F). Quadriceps were snap frozen in liquid nitrogen after dissection then stored at −80 °C.

Group 3 (M13-M18) animals were euthanized by intraperitoneal injection of pentobarbital/phenytoin (Euthasol^®^) followed by cervical dislocation. The intact carcasses were wrapped in foil and preserved by snap freezing in liquid nitrogen followed by storage at −80 °C. On the day of dissection, mouse carcasses were removed from the −80 °C freezer and thawed at room temperature for 15 to 20 minutes prior to dissection. Left lobes of livers and quadriceps were collected and snap frozen in liquid nitrogen then stored at −80 °C.

Group 4 (M19-M24) animals were euthanized by intraperitoneal injection of pentobarbital/phenytoin (Euthasol^®^) followed by cervical dislocation. The carcasses were sectioned into 3 sections, head, chest, and abdomen with tail removed and discarded and each part was submerged in RNA*later*™ solution (Thermo Fisher Scientific, Waltham, WA) and placed at 4 °C for 3 days to allow thorough permeation before being stored at −80 °C. On the day of dissection, mouse carcasses were removed from the −80 °C freezer and thawed at room temperature for 15 to 20 minutes prior to dissection. Left lobes of livers and quadriceps were collected and snap frozen in liquid nitrogen then stored at −80 °C. This group was used to simulate the procedure done in the RR-7 mission and to test if gene expression signals could be better preserved using an RNA-specific preservative.

Group 5 (M25-30) animals were euthanized by intraperitoneal injection of ketamine/xylazine (10mg/mL / 3mg/mL in 0.3mL PBS) followed by cervical dislocation. The intact carcasses were wrapped in foil and preserved by freezing on dry ice then stored at −80 °C. On the day of dissection, mouse carcasses were removed from the − 80 °C freezer and thawed at room temperature for 15 to 20 minutes prior to dissection. Left lobes of livers and quadriceps were collected and snap frozen in liquid nitrogen then stored at −80 °C. The euthanasia and preservation methods used in this group mimics the process used to generate RR-3 carcass liver samples (Smith et al., 2017). Ketamine/xylazine is currently the most common euthanasia method used in RR missions.

Group 6 (M31-M36) animals were euthanized by carbon dioxide inhalation followed by cervical dislocation. The carcasses were wrapped in foil and preserved by freezing on dry ice then stored at −80 °C. On the day of dissection, mouse carcasses were removed from the −80 °C freezer and thawed at room temperature for 15 to 20 minutes prior to dissection. Left lobes of livers and quadriceps were collected and snap frozen in liquid nitrogen then stored at −80 °C. This group represents the euthanasia method commonly used in terrestrial laboratories and was used to evaluate any effects on gene expression due to the drug-induced euthanasia methods that have been used in RR missions.

#### RNA Isolation

RNA was isolated from partial left liver lobe and partial quadriceps muscle tissues using the AllPrep DNA/RNA Mini Kit (Qiagen, Valencia, CA). Briefly, homogenization buffer was made by adding 1:100 volume ratio of beta-mercaptoethanol to RLT buffer and kept on ice until use. On average, 48.85 mg of left liver lobe and 20.08 mg of left or right quadriceps was cut using a sterile scalpel and immediately placed in 600 μL of the RLT buffer solution. Complete tissue dispersion was achieved using the hand-held Polytron PT1300D homogenizer with 5 mm standard dispersing aggregate by implementing 20 second homogenization periods at a speed of 20,000 RPM. Homogenized samples were centrifuged for 3 minutes at room temperature at 15,000 RPM to remove cell debris. The supernatant from each sample was used to isolate and purify RNA following the manufacturer’s protocol. RNA was treated with RNase-Free DNase (Qiagen, Valencia, CA) and eluted in 50 μL of RNase- and DNase-free H_2_O molecular grade water. RNA concentration was measured using Qubit 3.0 Fluorimeter (Thermo Fisher Scientific, Waltham, MA). RNA quality was assessed using the Agilent 2100 Bioanalyzer with the Agilent RNA 6000 Nano Kit or Agilent RNA 6000 Pico Kit (Agilent Technologies, Santa Clara, CA).

#### Library Preparation and Sequencing

Three microliters of Mix 1 or Mix 2 of ERCC ExFold RNA Spike-In (Thermo Fisher Scientific, Waltham, MA) at a dilution of 1:100 was added to 1.5 μg aliquots of RNA immediately after extraction. The two mixes were randomly distributed within the six experimental groups. In addition, Universal Human and Mouse Reference RNA samples (Agilent Technologies, Santa Clara, CA) were included as control samples in the library construction and sequencing.

Library construction was performed using 500 ng of ERCC-spiked total RNA with an average RIN of 7.8 for liver samples and 9.8 for quadriceps samples. Total RNA was depleted of the ribosomal fraction and libraries were constructed with TruSeq Stranded Total RNA with Ribo-Zero Gold kit (Illumina, San Diego, CA). Libraries were indexed using 1.5 μM Unique Dual Index adapters with Unique Molecular Identifiers (Integrated DNA Technologies, Coralville, IA) and 15 cycles of amplification were performed to reach desired library concentration. Library size was assessed with 4200 TapeStation (Agilent Technologies, Santa Clara, CA), targeting average size of 300 nt.

Libraries were multiplexed then quantified using Universal qPCR Master Mix (Kapa Biosystems, Wilmington, MA). The library pool was sequenced on an iSeq 100 (Illumina, San Diego, CA) to assess sample quality and pool balancing before large-scale sequencing. The final library pool (with 1% PhiX spike-in for instrument control) was sequenced on a NovaSeq 6000 using one S4 and one S2 Reagent Kit (Illumina, San Diego, CA), paired-end and 149 bp reads, targeting 60 million clusters for each experimental sample.

### RNA Sequencing Data Analysis

Raw RNA sequence data from the RR-1 NASA Validation flight liver (GLDS-48 and GLDS-168) samples, RR-1 CASIS liver samples (GLDS-47), and the ground-based studies designed to simulate and assess spaceflight euthanasia, carcass and tissue preservation, and/or storage protocols, GLDS-49, GLDS-235, and GLDS-236 were analyzed using the GeneLab standard RNAseq analysis pipeline. First, adapters were removed with Cutadapt (v2.3) (Martin, 2011) using the Trim Galore! (v0.6.2) wrapper. Raw and trimmed read quality were evaluated with FastQC (v0.11.8), and MultiQC (v1.7) was used to generate MultiQC reports. *Mus musculus* STAR and RSEM references were built using STAR (v2.7.1a) and RSEM (v1.3.1) (Li and Dewey, 2011), respectively, with ensembl genome version mm10-GRCm38 (Mus_musculus.GRCm38.dna.toplevel.fa), and the following gtf annotation file: Mus_musculus.GRCm38.96.gtf. Trimmed reads were aligned to the *Mus musculus* STAR reference with STAR (v2.7.1a) (Dobin et al., 2013) and aligned reads were quantified using RSEM (v1.3.1) (Li and Dewey, 2011).

The following samples were used for downstream analyses; GLDS-47: FLT and GC; GLDS-48: FLT-C, FLT-I, GC-C, and GC-I; GLDS-49: LN2-3d, LN2-1y, DI-1y; GLDS-168: RR1-FLT-wERCC and RR1-GC-wERCC; for GLDS-235 and GLDS-236, all samples indicated in Figure S4 and Tables S1 and S2 were included. For each GLDS dataset, quantification data from select samples were imported to R (v3.6.0) with tximport (v1.14.0) (Soneson et al., 2016) and normalized with DESeq2 (v1.26.0) (Love et al., 2014). All ERCC genes were removed prior to normalization. Differential expression analysis was performed in R (v3.6.0) using DESeq2 (v1.26.0) (Love et al., 2014); all groups were compared using the Wald test and the likelihood ratio test was used to generate the F statistic p-value. Gene annotations were assigned using the following Bioconductor and annotation packages: STRINGdb (Szklarczyk et al., 2019), PANTHER.db (Muller, 2017), and org.Mm.eg.db (Carlson, 2017).

### Transcript Integrity Analysis

The geneBody_coverage.py function from RSeQC (v3.0.1) (Wang et al., 2012) was used to assess coverage across the median 1000 expressed genes across all datasets. A transcript integrity metric was defined as the ratio between the coverage in a window corresponding to position 10-30% and 80-100% (relative to the entire gene length) and used in boxplots. To determine significance between groups the nonparametric Mann– Whitney U test was used as distributions were not normal.

### Gene Set Enrichment Analysis

Pairwise gene set enrichment analysis (GSEA) was performed on the normalized counts from select samples in GLDS-48, GLDS-168, GLDS-235, and GLDS-236 using the C5: Gene Ontology (GO) gene set (MSigDB v7.1) as described (Subramanian et al., 2005). All comparisons were performed using the phenotype permutation, except those involving GLDS-48 immediate samples, which used the gene set permutation due to low sample size. The ranked lists of genes were defined by the signal-to-noise metric, and the statistical significance were determined by 1000 permutations of the gene set. FDR ≤ 0.25 and FDR ≤ 0.05 were considered significant for comparisons using the phenotype and gene set permutations, respectively, according to the authors’ recommendation.

### Data Availability

All sequencing data is available at NASA GeneLab (www.genelab.nasa.gov) (Ray et al., 2019). polyA enrichment-based data from the RR-1 NASA Validation Mission samples are at GLDS-48 (Globus and Galazka, 2015). polyA enrichment-based data from the RR-1 CASIS samples are at GLDS-47 (Globus et al., 2015). Data from ground-based study of tissue storage conditions are at GLDS-49 (GeneLab, 2016). Ribodepletion-based resequencing data from the RR-1 NASA Validation Mission samples are at GLDS-168 (Galazka, 2018). Liver data from ground-based freezing study are at GLDS-235 (Galazka, 2019a). Quadriceps data from ground-based freezing study are at GLDS-236 (Galazka, 2019b).

## Acknowledgments

We thank Rebecca A. Klotz and Vandana Verma for dissection instructions; Candice G. T. Tahimic for help with IACUC protocol drafting; Samrawit G. Gebre for help with sample processing. This work was funded by the NASA Space Biology program within the NASA Science Mission Directorate’s (SMD) Biological and Physical Sciences (BPS) Division.

## Author Contributions

Formal analysis, AS.M.B, H.F, J.M.G; Investigation, K.C., R.B.C, O.M., S.L.P, V.B., M.T.D, Y.C., S.R.; Project administration, A.M.SB., S.V.C., J.M.G.; Visualization, A.M.SB, J.M.G; Writing—original draft, A.M.SB, S.V.C, J.M.G, S.L.P; Writing—review and editing, Y.C., A.M.SB, S.S.R., J.M.G., S.L.P; Conceptualization, S.L.P., J.M.G, S.V.C, S.S.R.; Data curation, A.M.SB., S.L.P., Y.C.

## Declaration of Interests

The authors declare no competing interests.

## Supplemental Information

### Supplemental Figure Legends

**Supplemental Figure 1.**
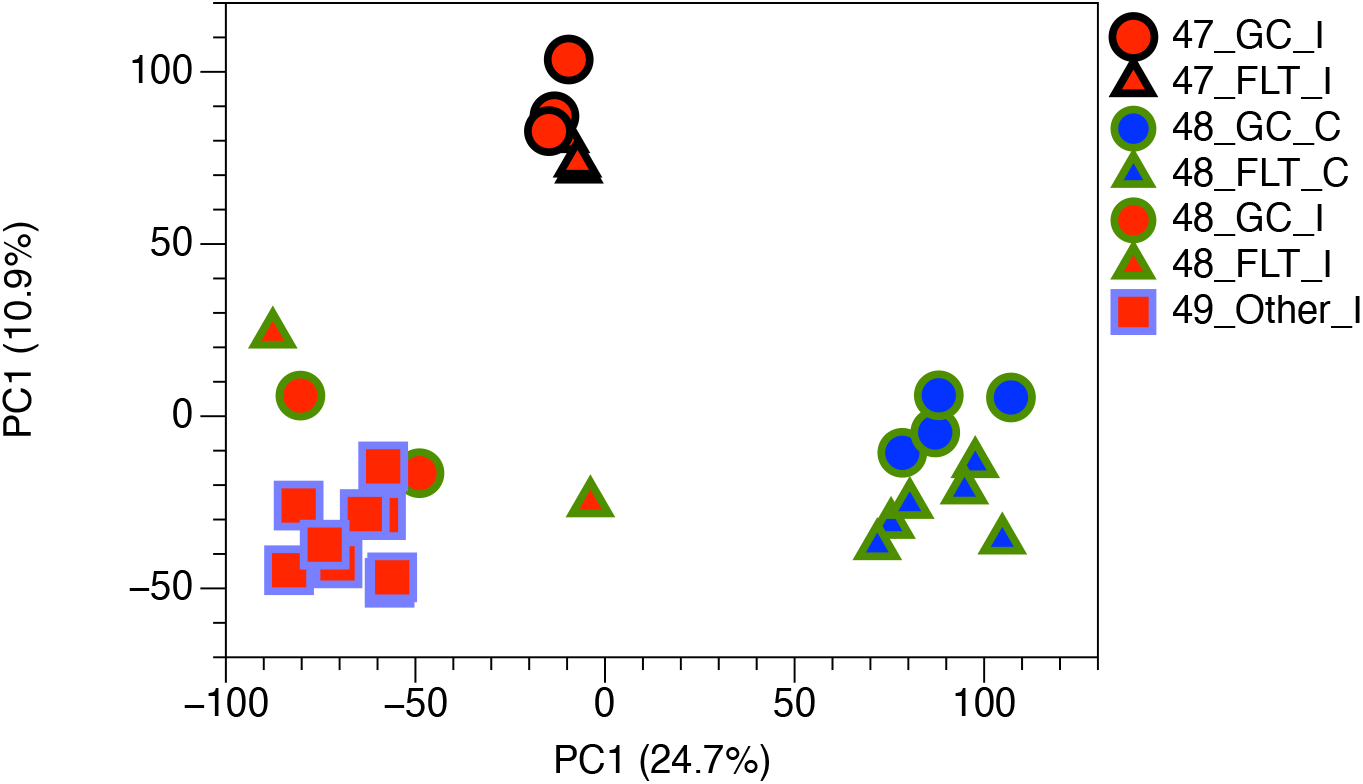
Carcass liver samples cluster together and away from immediate liver samples across datasets. Principal component analysis containing data from RR-1 NASA spaceflight (FLT) and respective ground control (GC) carcass (FLT_C_48 and GC_C_48) and immediate (FLT_I_48 and GC_I_48) samples (GLDS-48), RR-1 CASIS FLT (FLT_I_47) and GC (GC_I_47) immediate samples (GLDS-47), and samples from a ground-based study in which livers were dissected immediately after euthanasia then frozen on either dry ice (DI) or submerged in liquid nitrogen (LN2) then stored at −80 °C for either 3 days (LN2_I_3d_49) or 1 year (DI_I_1y_49 and LN2_I_1y_49) prior to processing. Related to Figure 1.

**Supplemental Figure 2.**
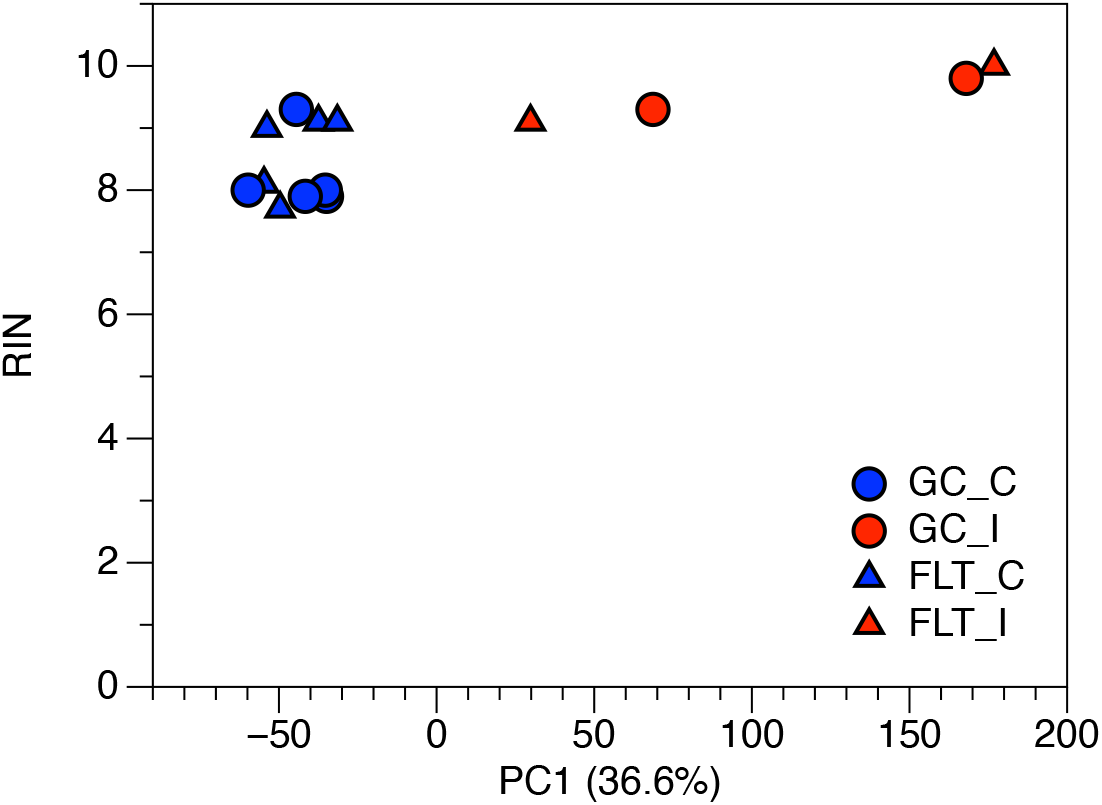
RN Integrity Number analysis of Carcass and Immediate Liver samples. RNA Integrity Numbers (RIN) for spaceflight (FLT) and ground control (GC) immediate (I) and carcass (C) samples plotted against principal component 1 (PC1) calculated from gene expression data. Related to Figures 1&3.

**Supplemental Figure 3.**
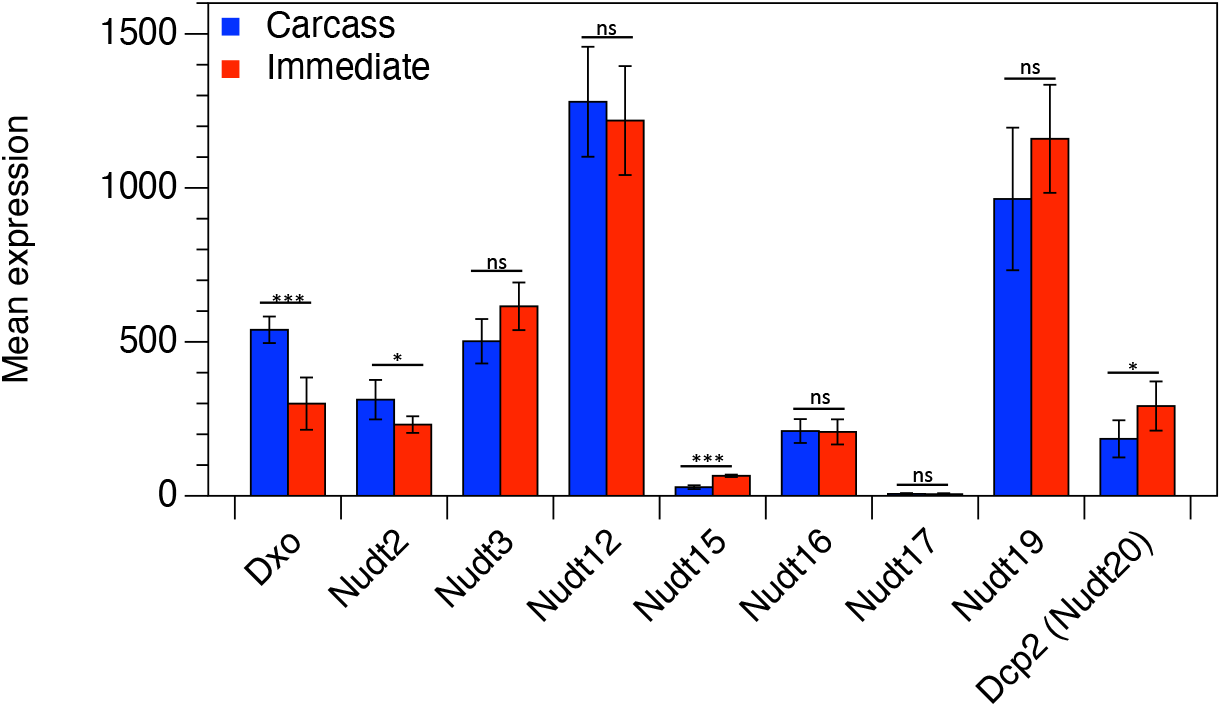
Expression of decapping enzymes. A) Average expression of decapping enzymes in Carcass (blue) and Immediate (red) groups. Error bars indicate standard deviation (* = adj. p < 0.05, ** = adj. p < 0.01, *** = adj. p < 0.001, ns = not significant, Wald test). Related to Figure 3.

**Supplemental Figure 4.**
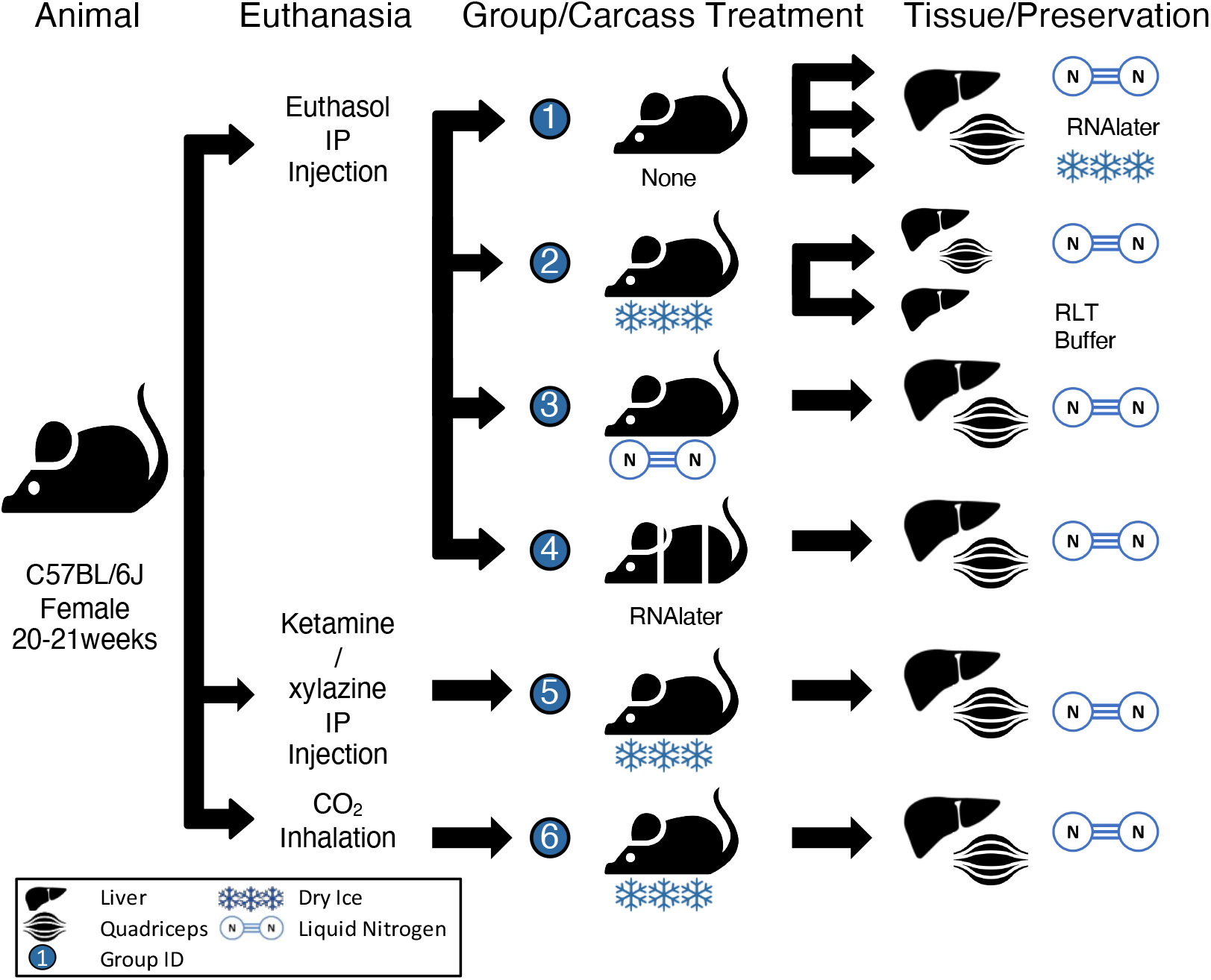
Freezing study workflow. Diagram of tissue preservation study to evaluate differences in indicated euthanasia, carcass and tissue preservation methods. Mice were euthanized with either pentobarbital/phenytoin (Euthasol^®^) or ketamine/xylazine injection, or CO2 inhalation. Intact carcasses were preserved by freezing in liquid nitrogen or on dry ice, or by segmentation (head, chest, abdomen) and immersion in an ammonium sulfate solution (RNA*later*™). Carcasses were then thawed and livers and quadriceps dissected and preserved in liquid nitrogen or guanidinium thiocyanate solution (Qiagen^®^ RLT buffer). Alternatively, livers and quadriceps were dissected immediately and preserved by freezing in liquid nitrogen or on dry ice, or by immersion in an ammonium sulfate solution (RNA*later*™). Related to Figure 5 and Tables 1&2.

**Supplemental Figure 5.**
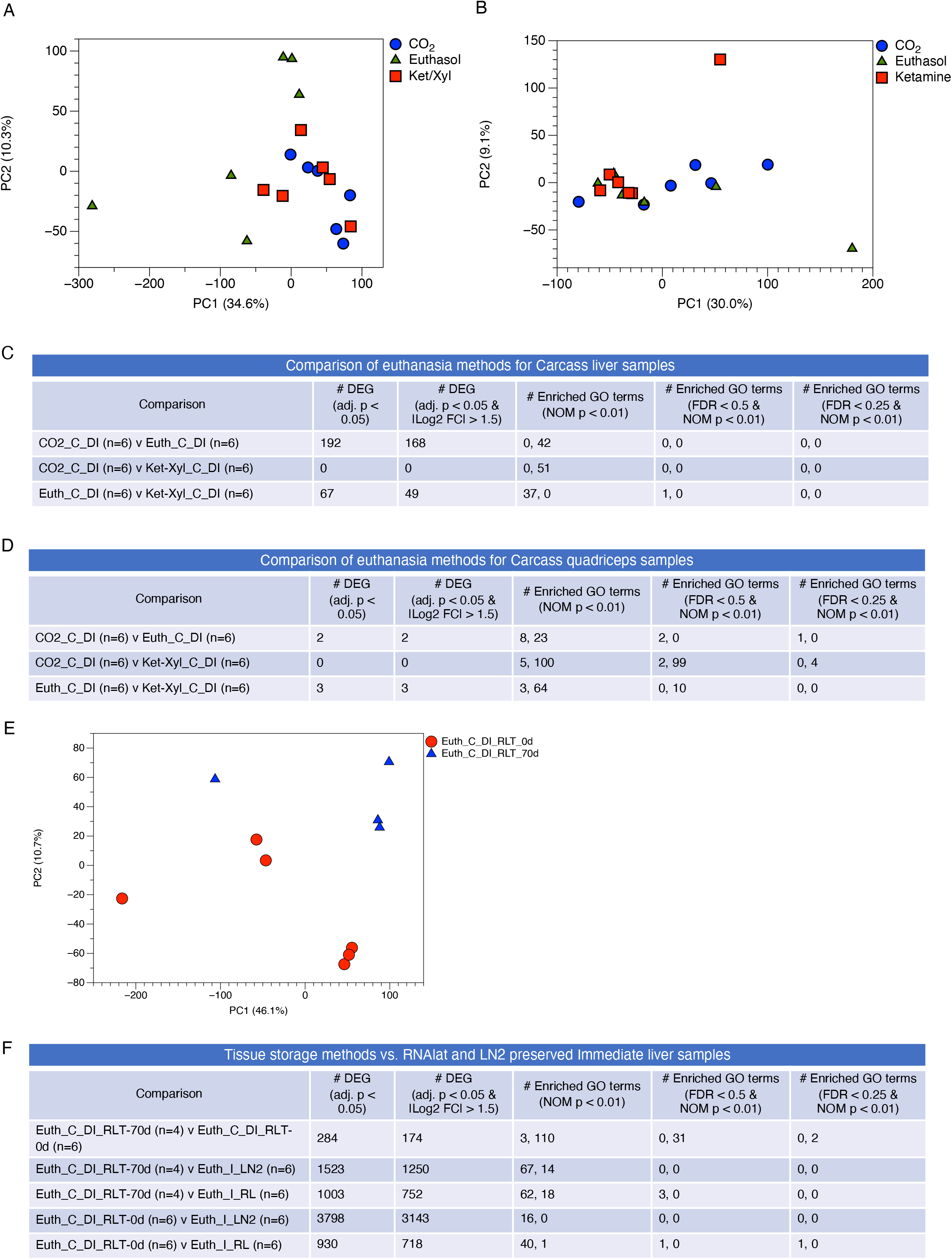
Comparison of gene expression and gene ontology in liver and quadriceps samples derived from mice euthanized with different methods. Liver and quadriceps samples dissected from partially thawed frozen carcasses of mice that were euthanized with pentobarbital/phenytoin (Euthasol^®^), ketamine/xylazine, or carbon dioxide inhalation were evaluated for global gene expression differences via principal component analysis (A, liver and B, quadriceps), and the number of differentially expressed genes (DEG) and enriched gene ontology (GO) terms identified by Gene Set Enrichment Analysis via pairwise comparisons (phenotype permutation) (C, liver and D, quadriceps). Liver samples dissected from partially thawed frozen carcasses of mice that were euthanized with pentobarbital/phenytoin (Euthasol^®^), then snap frozen in liquid nitrogen and stored at −80 °C or homogenized in RLT buffer then stored for 70 d at −80 °C were evaluated for E) global gene expression differences via principal component analysis, and F) the number of differentially expressed genes (DEG) and enriched gene ontology (GO) terms identified by Gene Set Enrichment Analysis (phenotype permutation) via pairwise comparisons with immediate samples preserved in liquid nitrogen or RNAlater. For GO terms, the number on the left corresponds to the group to the left of the ‘vs.’, and number on the right corresponds to the group to the right of the ‘vs.’ in the “Comparison” column. n numbers, p values, log2 fold changes, and FDR values are indicated. Euth=euthanasia by pentobarbital/phenytoin (Euthasol^®^), Ket-Xyl= euthanasia by ketamine/xylazine, CO2=euthanasia by carbon dioxide inhalation, C=tissue dissected from frozen carcass that has been partially thawed, I=tissue dissected immediately after euthanasia, DI=dry ice, LN2=liquid nitrogen, RL=RNA*later*™. Liver and quadriceps data are from GLDS-235 and GLDS-236, respectively. Related to Figure 5 and Tables 1&2.

**Supplemental Figure 6.**
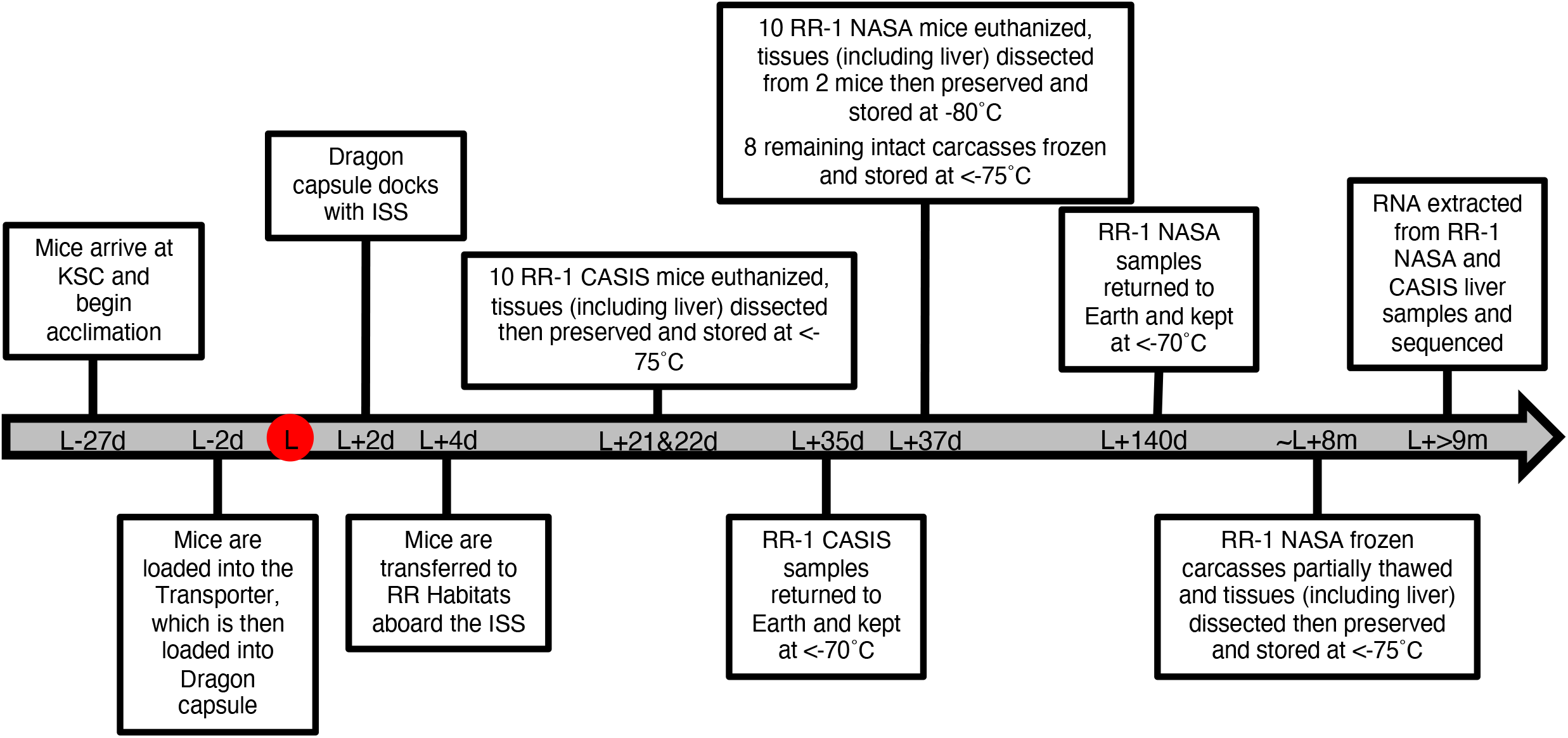
RR-1 mission timeline. Timeline indicating major events in the Rodent Research-1 (RR-1) mission relative to the launch date (L). A minus sign (-) indicates time in days (d) before launch and a plus sign (+) indicates time in days (d) or months (m) after launch. Age-matched ground control animals were processed on similar timeline but on a 4-day delay to mimic spaceflight conditions. KSC = Kennedy Space Center; ISS = International Space Station; CASIS = Center for the Advancement of Science in Space; NASA = National Aeronautics and Space Administration. Related to Figures 1-4.

**Supplemental Figure 7.**
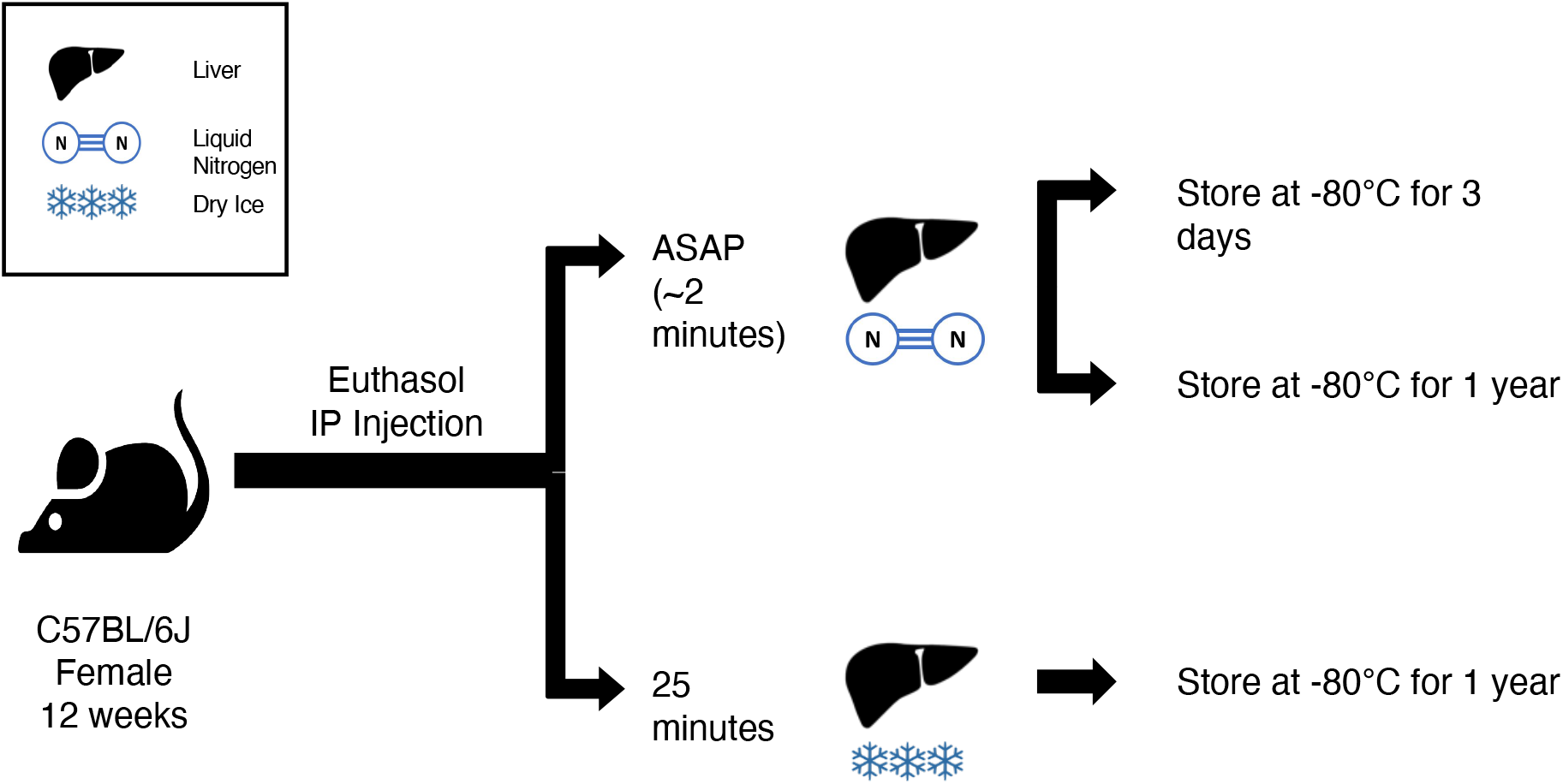
GLDS-49 Experimental Design. Diagram of the ground-based preservation study comparing livers collected using standard laboratory protocols with livers collected from simulated spaceflight dissection flow and storage methods. Liver samples from twelve-week old female C57BL/6J mice were either snap frozen (n=3), snap frozen after a 25 min delay and stored for 3 days (n=3), or snap frozen after a 25 min delay and stored for 1 year (n=3). RNA-seq data were then generated using a polyA enrichment protocol. Related to Figure 1.

### Supplementary Tables

**Table S1.** Liver samples analyzed in the ground-based tissue preservation study. Related to Figure 5A&B and Table 1.

| Group | Euthanasia | Carcass preservation | Tissue preservation | Mouse ID |
| --- | --- | --- | --- | --- |
| 1 | Pentobarbital/phenytoin | None | Liquid nitrogen n=6 | M1-M6 |
|  |  |  | RNAlater™ n=6 |  |
|  |  |  | Dry ice n=6 |  |
| 2 | Pentobarbital/phenytoin | Dry ice | Liquid nitrogen n=6 | M7-M12 |
|  |  |  | RLT buffer 70d at -80 °C n=4 |  |
| 3 | Pentobarbital/phenytoin | Liquid nitrogen | Liquid nitrogen n=5 | M13-M18 |
| 4 | Pentobarbital/phenytoin | RNAlater™<sup>a</sup> | Liquid nitrogen n=5 | M19-M24 |
| 5 | Ketamine/xylazine | Dry ice | Liquid nitrogen n=6 | M25-M30 |
| 6 | Carbon dioxide | Dry ice | Liquid nitrogen n=6 | M31-M36 |

**Table S2.**
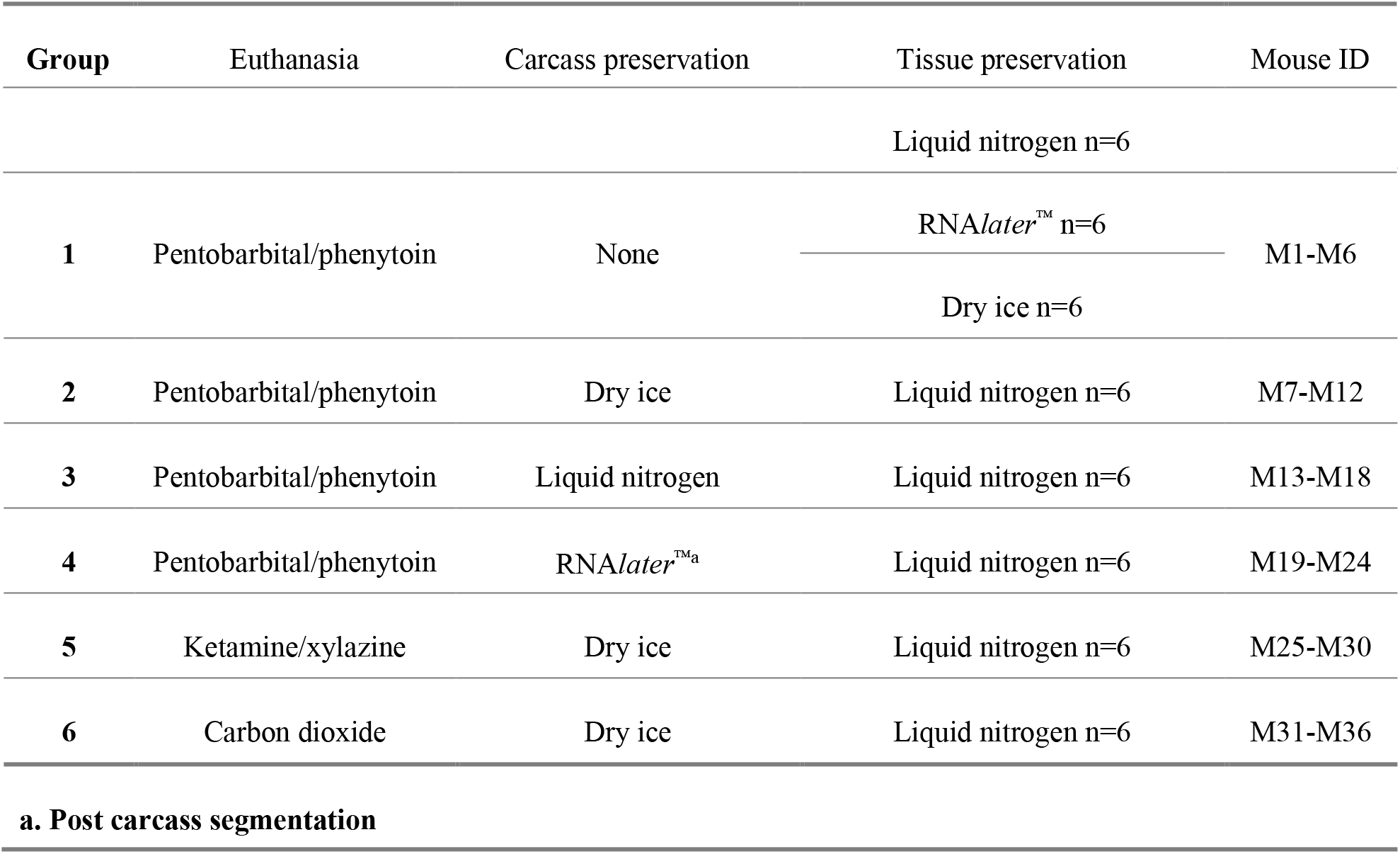
Quadriceps samples analyzed in the ground-based tissue preservation study. Related to Figure 5C&D and Table 2.

**Table S3.** Comparisons of immediate preservation methods on gene expression in livers. The number of differentially expressed genes (DEG) and enriched gene ontology (GO) terms identified by Gene Set Enrichment Analysis (phenotype permutation) were evaluated pairwise in liver samples from different immediate preservation methods. For GO terms, the first number corresponds to the group to the left of the ‘vs.’, and second number corresponds to the group to the right of the ‘vs.’ in the “Comparison” column. n numbers, p values, log2 fold changes, and FDR values are indicated. Euth=euthanasia by pentobarbital/phenytoin, I=tissue dissected immediately after euthanasia, DI=dry ice, LN2=liquid nitrogen, RL=RNA*later*™. Data are from GLDS-235. Related to Figure 5A&B and Table 1.

| Comparison | # DEG<br>(adj. p < 0.05) | # DEG<br>(adj. p < 0.05 & \|Log2 FC\| > 1.5) | # Enriched GO terms<br>(NOM p < 0.01) | # Enriched GO terms<br>(FDR < 0.5 & NOM p < 0.01) | # Enriched GO terms<br>(FDR < 0.25 & NOM p < 0.01) |
| --- | --- | --- | --- | --- | --- |
| Euth_I_DI (n=6) vs. Euth_I_LN2 (n=6) | 16 | 16 | 31, 6 | 7, 0 | 0, 0 |
| Euth_I_DI (n=6) vs. Euth_I_RL (n=6) | 0 | 0 | 14, 8 | 0, 0 | 0, 0 |
| Euth_I_LN2 (n=6) vs. Euth_I_RL (n=6) | 14 | 14 | 3, 15 | 0, 0 | 0, 0 |

**Table S4.** Comparisons of carcass preservation methods to immediate RNA*later*™ method on gene expression in livers. The number of differentially expressed genes (DEG) and enriched gene ontology (GO) terms identified by Gene Set Enrichment Analysis (phenotype permutation) were evaluated pairwise in liver samples from different carcass preservation methods compared with immediate samples preserved in RNA*later*™. For GO terms, the first number corresponds to the group to the left of the ‘vs.’, and the second number corresponds to the group to the right of the ‘vs.’ in the “Comparison” column. n numbers, p values, log2 fold changes, and FDR values are indicated. Euth=euthanasia by pentobarbital/phenytoin, I=tissue dissected immediately after euthanasia, C=tissue dissected from frozen carcass that has been partially thawed, DI=dry ice, LN2=liquid nitrogen, RL= RNA*later*™. Data are from GLDS-235. Related to Figure 5A&B and Table 1.

| Comparison | # DEG<br>(adj. p < 0.05) | # DEG<br>(adj. p < 0.05 &<br>\|Log2 FC\| > 1.5) | # Enriched GO<br>terms<br>(NOM p < 0.01) | # Enriched GO<br>terms<br>(FDR < 0.5 &<br>NOM p < 0.01) | # Enriched GO<br>terms<br>(FDR < 0.25 &<br>NOM p < 0.01) |
| --- | --- | --- | --- | --- | --- |
| Euth_C_DI (n=6)<br>vs. Euth_I_RL<br>(n=6) | 930 | 718 | 40, 1 | 1, 0 | 1, 0 |
| Euth_C_LN2<br>(n=5) vs.<br>Euth_I_RL (n=6) | 197 | 118 | 90, 0 | 2, 0 | 0, 0 |
| Euth_C_RL<br>(n=5) vs.<br>Euth_I_RL (n=6) | 131 | 123 | 65, 0 | 0, 0 | 0, 0 |

**Table S5.** Comparisons of immediate preservation methods on gene expression in quadriceps. The number of differentially expressed genes (DEG) and enriched gene ontology (GO) terms identified by Gene Set Enrichment Analysis (phenotype permutation) were evaluated pairwise in quadriceps samples from different immediate preservation methods. For GO terms, number on the left corresponds to the group to the left of the ‘vs.’, and number on the right corresponds to the group to the right of the ‘vs.’ in the “Comparison” column. n numbers, p values, log2 fold changes, and FDR values are indicated. Euth=euthanasia by pentobarbital/phenytoin, I=tissue dissected immediately after euthanasia, DI=dry ice, LN2=liquid nitrogen, RL= RNA*later*™. Data are from GLDS-236. Related to Figure 5C&D and Table 2.

| Comparison | # DEG<br>(adj. p < 0.05) | # DEG<br>(adj. p < 0.05 &<br>\|Log2 FC\| > 1.5) | # Enriched GO<br>terms<br>(NOM p < 0.01) | # Enriched GO<br>terms<br>(FDR < 0.5 &<br>NOM p < 0.01) | # Enriched GO<br>terms<br>(FDR < 0.25 &<br>NOM p < 0.01) |
| --- | --- | --- | --- | --- | --- |
| **Euth_I_DI (n=6)**<br>**vs. Euth_I_LN2**<br>**(n=6)** | 2 | 1 | 2, 51 | 1, 2 | 0, 2 |
| **Euth_I_DI (n=6)**<br>**vs. Euth_I_RL**<br>**(n=6)** | 14 | 9 | 25, 26 | 6, 0 | 2, 0 |
| **Euth_I_LN2**<br>**(n=6) vs.**<br>**Euth_I_RL (n=6)** | 0 | 0 | 66, 3 | 51, 0 | 0, 0 |

**Table S6.**
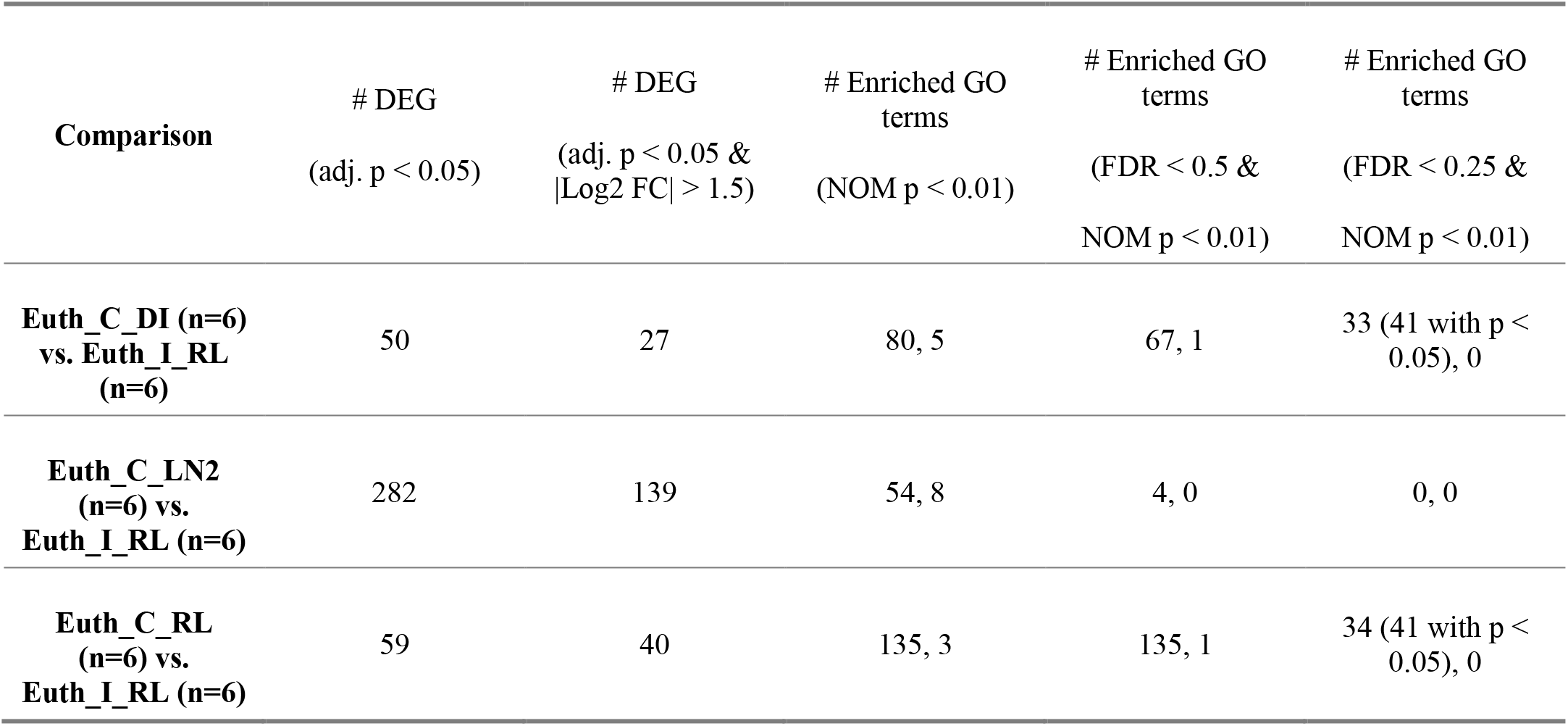
Comparisons of carcass preservation methods to immediate RNA*later*™ method on gene expression in quadriceps. The number of differentially expressed genes (DEG) and enriched gene ontology (GO) terms identified by Gene Set Enrichment Analysis (phenotype permutation) were evaluated pairwise in quadriceps samples from different carcass preservation methods compared with immediate samples preserved in RNA*later*™. For GO terms, number on the left corresponds to the group to the left of the ‘vs.’, and number on the right corresponds to the group to the right of the ‘vs.’ in the “Comparison” column. n numbers, p values, log2 fold changes, and FDR values are indicated. Euth=euthanasia by pentobarbital/phenytoin, I=tissue dissected immediately after euthanasia, C=tissue dissected from frozen carcass that has been partially thawed, DI=dry ice, LN2=liquid nitrogen, RL= RNA*later*™. Data are from GLDS-236. Related to Figure 5C&D and Table 2.

| **Comparison** | **# DEG**<br>**(adj. p < 0.05)** | **# DEG**<br>**(adj. p < 0.05 &**<br>**\|Log2 FC\| > 1.5)** | **# Enriched GO**<br>**terms**<br>**(NOM p < 0.01)** | **# Enriched GO**<br>**terms**<br>**(FDR < 0.5 &**<br>**NOM p < 0.01)** | **# Enriched GO**<br>**terms**<br>**(FDR < 0.25 &**<br>**NOM p < 0.01)** |
| --- | --- | --- | --- | --- | --- |
| **Euth_C_DI (n=6)**<br>**vs. Euth_I_RL**<br>**(n=6)** | 50 | 27 | 80, 5 | 67, 1 | 33 (41 with p < 0.05), 0 |
| **Euth_C_LN2**<br>**(n=6) vs.**<br>**Euth_I_RL (n=6)** | 282 | 139 | 54, 8 | 4, 0 | 0, 0 |
| **Euth_C_RL**<br>**(n=6) vs.**<br>**Euth_I_RL (n=6)** | 59 | 40 | 135, 3 | 135, 1 | 34 (41 with p < 0.05), 0 |

### Supplemental Materials and Methods

#### GLDS-49 Ground-based Freezing Study

To compare standard laboratory protocols for tissue freezing and storage with a spaceflight timeline-simulated liver dissection and long-term storage, liver samples from twelve-week old female C57BL/6J mice (Jackson laboratories, Bar harbor, ME) were either immediately snap frozen (in liquid nitrogen), snap frozen after a 25 min delay and stored for 3 days, or snap frozen after a 25 min delay and stored for 1 year (Figure S7).

#### Sample Collection

The liver tissues of twelve-week-old C57BL/6J mice (Jackson laboratories, Bar harbor, ME) were received from the Rodent Research project collected in a ground-based preservation and storage study (Choi et al., 2016). Three groups of livers were included: 1) Liver tissues dissected and frozen on dry ice 25 min after euthanizing with pentobarbital/phenytoin (Euthasol^®^) followed by cervical dislocation. At the time of RNA extraction, the liver tissues had been stored at −80 °C for around 1-year (DI_I_1y_49); 2) Liver tissues dissected 3 min after euthanizing with pentobarbital/phenytoin (Euthasol^®^) followed by cervical dislocation and snap-freezing in liquid nitrogen. At the time of RNA extraction, the liver tissues had been stored at −80 °C for around 1-year (LN_I_1y_49); 3) Liver tissues dissected 3 minutes after euthanizing with pentobarbital/phenytoin (Euthasol^®^) followed by cervical dislocation and snap-freezing in liquid nitrogen. At the time of RNA extraction, the liver tissues had been stored at − 80 °C for only 3 days (LN_I_3d_49). This group served as a positive control for delayed dissection and long-term storage.

#### RNA Isolation

RNA was isolated using the AllPrep DNA/RNA Mini Kit (Qiagen, Valencia, CA) following the manufacturer’s protocol. Briefly, homogenization buffer was made by adding 1:100 volume ratio of beta-mercaptoethanol to RLT buffer and kept on ice until use. Approximately 30 mg of tissue was cut using a sterile scalpel and immediately placed in 800 µL of the RLT buffer solution. Each sample was then homogenized for approximately 20 seconds at 21,000 RPM using a Polytron PT1300D handheld homogenizer with a 5 mm standard dispersing aggregate tip (Kinematica, Bohemia, NY). Homogenates were centrifuged for 3 minutes at room temperature at 15,000 RPM to remove cell debris. The supernatant from each sample was used to isolate and purify RNA following the manufacturer’s protocol including on-column DNase treatment with RNase-free DNase (Qiagen, Valencia, CA). RNA was eluted twice per sample in 30 µL RNase- and DNase-free H2O per elution. Concentration and absorbance ratios were measured using the NanoDrop 2000 spectrophotometer (Thermo Fisher Scientific, Waltham, MA). RNA quality was assessed using the Agilent 2100 Bioanalyzer with the Agilent RNA 6000 Nano Kit (Agilent Technologies, Santa Clara, CA).

#### Library Preparation and RNA-Sequencing

Samples with RNA Integrity Number (RIN) of 9 or above were sent to the University of California (UC), Davis Genome Center where the libraries were constructed and RNA-sequencing was performed. Libraries were generated using the Illumina TruSeq Stranded RNA library prep kit (Illumina, San Diego, CA) after polyA selection, and sequencing was done with 50 bp single end reads on the Illumina HiSeq 3000 platform.

## Notes

### Competing Interest Statement

The authors have declared no competing interest.

